# CRISPR screens reveal genetic determinants of PARP inhibitor sensitivity and resistance in prostate cancer

**DOI:** 10.1101/2022.04.13.488115

**Authors:** Takuya Tsujino, Tomoaki Takai, Kunihiko Hinohara, Fu Gui, Takeshi Tsutsumi, Xiao Bai, Chenkui Miao, Chao Feng, Bin Gui, Zsofia Sztupinszki, Antoine Simoneau, Ning Xie, Ladan Fazli, Xuesen Dong, Haruhito Azuma, Atish D. Choudhury, Kent W. Mouw, Zoltan Szallasi, Lee Zou, Adam S. Kibel, Li Jia

**Author notes:** These authors contributed equally to this article (Takuya Tsujino, Tomoaki Takai, Kunihiko Hinohara).

## Abstract

Prostate cancer (PCa) harboring BRCA1/2 mutations is often exquisitely sensitive to PARP inhibition. However, genomic alterations in other DNA damage response genes have not been consistently predictive of clinical response to PARP inhibitors (PARPis). Here, we perform genome-wide CRISPR-Cas9 knockout screens in BRCA1/2-proficient PCa cell lines and identify novel genes whose loss has a profound impact on PARPi sensitivity and resistance. Specifically, MMS22L deletion, frequently observed (up to 14%) in PCa, renders cells hypersensitive to PARPis by disrupting RAD51 loading required for homologous recombination repair, although this response is TP53-dependent. Unexpectedly, loss of CHEK2 confers resistance rather than sensitivity to PARPis in PCa cells through increased expression of BRCA2, a target of CHEK2-TP53-E2F7-mediated transcriptional repression. Combined PARP and ATR inhibition overcomes PARPi resistance caused by CHEK2 loss. Our findings may inform the use of PARPis beyond BRCA1/2-deficient tumors and support reevaluation of currently used biomarkers for PARPi treatment in PCa.

## INTRODUCTION

Despite treatment advances, metastatic castration-resistant prostate cancer (mCRPC) remains a lethal disease. Genomic sequencing studies reveal that approximately 90% of mCRPC patients carry clinically actionable genomic alterations^1^. Alterations in genes involved in the DNA damage response (DDR) are among the most common genetic events. Approximately 10% of primary and 25% of metastatic PCa patients have an alteration in at least one gene involved in DDR^2^, which represents a potential therapeutic vulnerability. In particular, defects in homologous recombination repair (HRR) render cells highly sensitive to inhibition of Poly (ADP-ribose) polymerase (PARP). As a targeted therapy, PARP inhibitors (PARPis) prevent PARP1 and PARP2 from repairing DNA single-strand breaks (SSBs) and leads to stalled and collapsed replication forks by trapping PARP1/2 on the DNA breaks^3^. Subsequently, SSBs are converted to double-strand breaks (DSBs) that HRR-deficient cells cannot repair effectively, leading to overwhelming DNA damage, cell cycle arrest, and cell death. The BRCA1 and BRCA2 genes encode proteins essential for HRR, a pathway that repair DNA DSBs. PARPis selectively induce synthetic lethality in cancer cells harboring mutations in the BRCA1/2 genes^4,5^

Several PARPis are presently under clinical investigation in PCa, including olaparib, rucaparib, niraparib, and talazoparib, as a single agent. Two of them (olaparib and rucaparib) have been approved by the U.S. Food and Drug Administration (FDA) for the treatment of mCRPC patients with deleterious germline and/or somatic mutations in BRCA1/2, and the olaparib indication includes mutations in 12 additional HRR genes^6–11^. It is clear that BRCA1/2-mutant tumors exhibit high sensitivity and improved outcome to PARP inhibition based on results from these trials, as measured by standard radiographic criteria, 50% decrease in prostate-specific antigen (PSA), circulating tumor-cell counts, and progression-free survival or overall survival. However, whether and to what extent PARPis can be used to treat tumors with non-BRCA1/2 alterations remains controversial after gene-by-gene analysis. Furthermore, HRR-deficient tumors are not always sensitive to PARPis. Both intrinsic and acquired resistance to PARP inhibition represents a formidable clinical problem. Therefore, one of the major barriers to effective treatment using PARPis is how to distinguish patients who may and may not benefit from PARP inhibition.

The genome-wide CRISPR-Cas9 (clustered regularly interspersed short palindromic repeat-CRISPR associated nuclease 9) knockout (KO) screen is a powerful and unbiased approach to identify genes that, when deleted, confer PARPi sensitivity or resistance. Previous studies using CRISPR screens in non-PCa cell lines have identified genes whose loss impacts PARPi response^12–14^. In this study, we carried out CRISPR screens in four BRCA1/2-proficient PCa cell lines with the goal of expanding the use of PARPis beyond BRCA1/2 mutations and finding new synthetically lethal interactions. Our screens led to the identification of MMS22L, a frequently deleted HRR gene in PCa, as a new predictive biomarker for PARPis. Unexpectedly, we found that loss of CHEK2 caused PARPi resistance, which we explored a therapeutic approach through ATR inhibition to overcome in pre-clinical models.

## RESULTS

### Genome-wide CRISPR KO screens identify genes that modulate PARPi response in PCa cells

To identify genes whose loss increases or decreases the sensitivity of PCa cells to PARP inhibition, we performed genome-wide CRISPR KO screens in the PCa LNCaP, C4-2B, 22Rv1 and DU145 cells in the presence of olaparib or DMSO vehicle. These four cell lines reflect different aspects of PCa progression to castration resistance and have no deleterious biallelic mutations in BRCA1/2 and other canonical HRR genes determined by whole exome sequencing (Supplementary Table 1). LNCaP and C4-2B cells are relatively more sensitive to olaparib in contrast to 22Rv1 and DU145 cells (Supplementary Fig. 1), but less sensitive when compared to BRCA1-null ovarian cancer UWB1 cells. Cells were transduced with the lentiviral-based CRISPR-Cas9 KO libraries, targeting over 18,000 protein-coding genes as previously described^15,16^, followed by 28 days of treatment with olaparib or DMSO (Fig. 1a; Supplementary Fig. 2a,b). The abundance of single guide RNA (sgRNA) for each gene was assessed by β-score and the differential β-score was further calculated using the MAGeCKFlute pipeline by comparing olaparib treatment to DMSO treatment^17,18^. We identified 216, 243, 153, and 211 negatively selected genes (with a stringent cutoff of *p*-value < 0.01, σ = 2.58), loss of which sensitizes cells to olaparib in LNCaP, C4-2B, 22Rv1 and DU145 cells, respectively (Fig. 1b; Supplementary Table 2). There were 67 genes shared by at least two cell lines, considered as common hits (Supplementary Table 3). Gene Ontology (GO) analyses revealed that these common hits were enriched for DNA repair and replication functions (Fig. 1c). Further analysis of each individual cell line showed that the mitochondrial complex I assembly was the most enriched function in DU145 (Fig. 1d; Supplementary Fig. 3). Dysfunction of mitochondrial complex I causes a decline of NAD^+^ and ATP supply required for PARP-mediated DNA repair^19^. The cell-type specific mechanism may provide a unique therapeutic vulnerability.

**Fig. 1.**
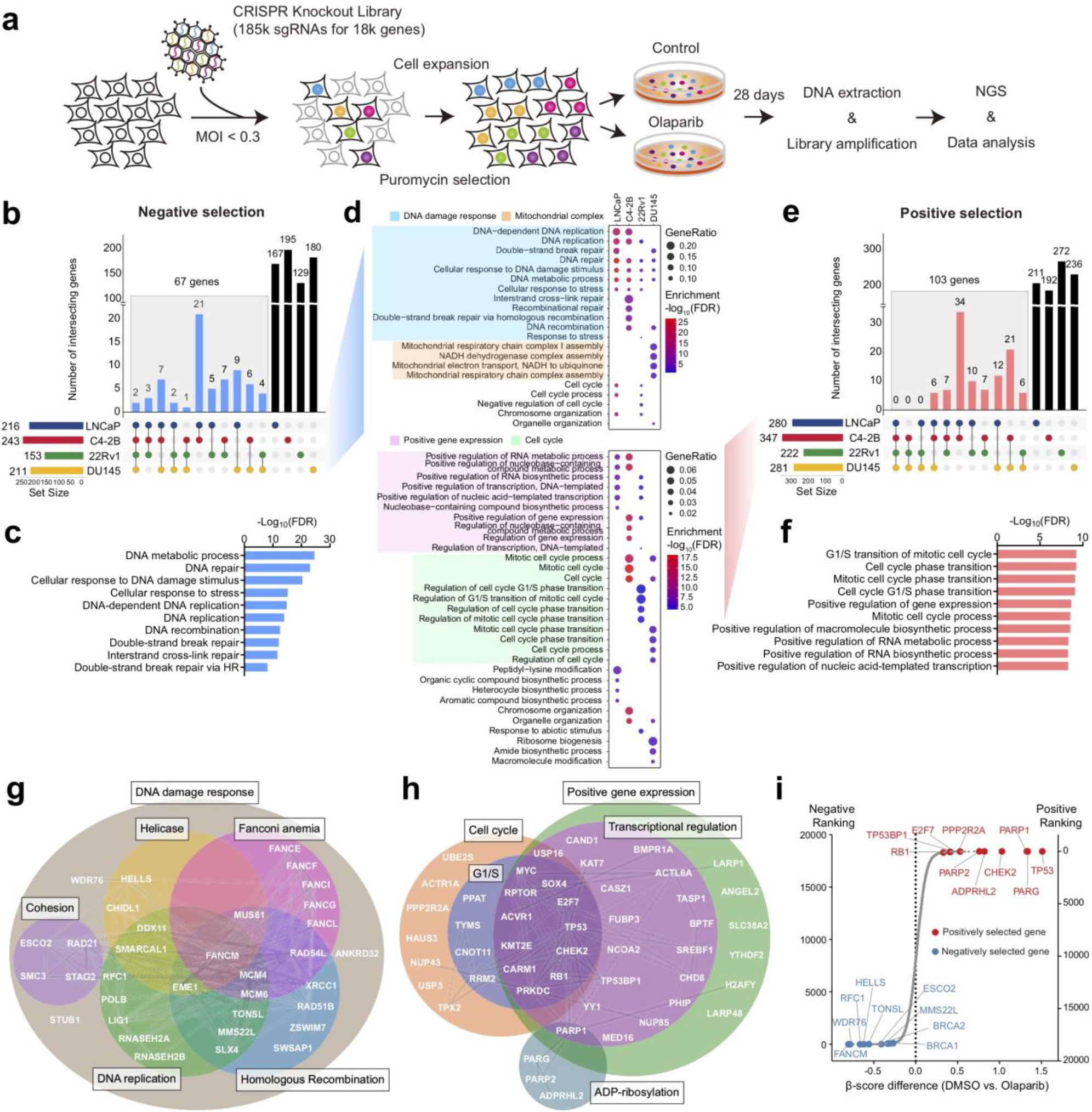
CRISPR screens identify genes that modulate PARPi response in PCa cells. **a,** Schematic of genome-wide CRISPR/Cas9 screens. **b,** UpSet plot^96^ of negatively selected genes in four PCa cell lines as indicated. Blue bars indicate the number of common hits in at least two screens. **c,** Top GO terms enriched in 67 common hits from negative selection. **d,** Top GO terms enriched in negatively (upper panel) and positively (lower panel) selected genes in each individual cell line. **e,** UpSet plot of positively selected genes in four PCa cell lines as indicated. Red bars indicate the number of common hits in at least two screens. **f,** Top GO terms enriched in 103 common hits from positive selection. **g,** The networks of common hits from negative selection, grouped according to their roles in specific pathways, and their genetic and physical interactions (gray lines) based on STRING analyses. **h,** The networks of common hits from positive selection as described in (**g**). **i,** Top-ranked genes in CRISPR screens determined by comparing olaparib treatment to DMSO treatment. Genes are ranked by the average of differential β-scores from all four cell lines. Negatively and positively selected genes are marked in blue and red, respectively.

We identified 280, 347, 222, and 281 positively selected genes (*p* < 0.01, σ = 2.58), loss of which renders cells resistant to olaparib in LNCaP, C4-2B, 22Rv1, and DU145 cells, respectively (Fig. 1e; Supplementary Table 4). Given that cells were cultured in the presence of olaparib for only 28 days, these genes are more likely related to intrinsic resistance rather than acquired resistance arising after prolonged treatment with PARPis. There were 103 genes shared in at least two cell lines (Supplementary Table 5). These common hits were enriched in cell cycle phase transition and positive regulation of transcription (Fig. 1f). Analyses of positively selected genes in each individual cell line showed that cell cycle genes are largely enriched in CRPC C4-2B, 22Rv1, and DU145 cells, but not in androgen-dependent LNCaP cells (Fig. 1d; Supplementary Fig. 3), reflecting the role of cell cycle transition in mediating PARPi resistance in CRPC^20^.

To further dissect the identified genes implicated in PARPi sensitivity and resistance, we analyzed the common hits using the database in STRING protein-protein association network^21^. We found that 50% of the common negatively selected genes could be assigned to well-defined functions in DDR, including HRR, DNA replication, Fanconi anemia, helicase, and cohesion (Fig. 1g). In contrast, approximately 50% of the common positively selected genes were involved specifically in cell cycle, ADP-ribosylation, transcriptional regulation (Fig. 1h). Finally, we ranked all genes based on the average of differential β-scores from all four cell lines. Representative top-ranked negatively and positively selected genes are shown in Fig. 1i, including ones studied in this work. Notably, the BRCA1 and BRCA2 genes were highly ranked in negative selection but reached the cutoff only in C4-2B and DU145 cells, respectively. This is likely because acute inactivation of BRCA1 and BRCA2 is lethal without adaptive mechanisms in cells^22–24^, as reflected by the depletion of sgRNAs targeting BRCA1/2 genes even in the absence of olaparib treatment in most cases, thereby failing to reach statistically significant depletion in the olaparib condition (Supplementary Fig. 4). Together, our screen results provide a global view of genetic determinants of PARPi response in PCa.

### Validation of negatively selected genes related to PARPi sensitivity

Next, we examined the alteration frequency (deletion and mutation) of the 67 common negatively selected genes in the TCGA (primary tumors) and SU2C/PCF (metastatic tumors) cohorts (Fig. 2a). We selected ten genes, based on their alteration frequency as well as their important functions in DDR, and individually deleted these genes in C4-2B cells (Supplementary Fig. 5). We observed markedly increased sensitivity to olaparib following deletion (Fig. 2b). Among these genes, HELLS and WDR76 are the only two hits in all four cell lines. HELLS is a helicase and chromatin remodeling enzyme that promotes the initiation of HRR and contributes to DSB repair^25^, while WDR76 is a specific reader of 5-hydroxymethylcytosine (5hmC) with uncharacterized functions in DNA repair^26,27^. Indeed, WDR76 has a large interaction network, including HELLS and PARP1^28^, supporting synthetically lethal interaction with PARP inhibition. RNASEH2B and MMS22L are located on chromosomes 13q14 and 6q16, respectively, which are two genomic regions frequently deleted in PCa^29–35^. While loss of RNASEH2B might cause ribonucleotide excision repair deficiency and PARP-trapping lesions as previously reported^13^, we recently reported that co-loss of RB1, a closely located tumor suppressor gene, was antagonistic in PCa cells^36^. MMS22L is known to form an obligate heterodimer complex with TONSL, which reads histone H4 unmethylated lysine 20^37^ and promotes HRR of stalled or collapsed replication forks and attendant DSBs by aiding RAD51 loading^38–40^. Cells with MMS22L deletion are highly sensitive to the topoisomerase I inhibitor camptothecin. However, the impact of MMS22L loss on PARP inhibition has not been studied.

**Fig. 2.**
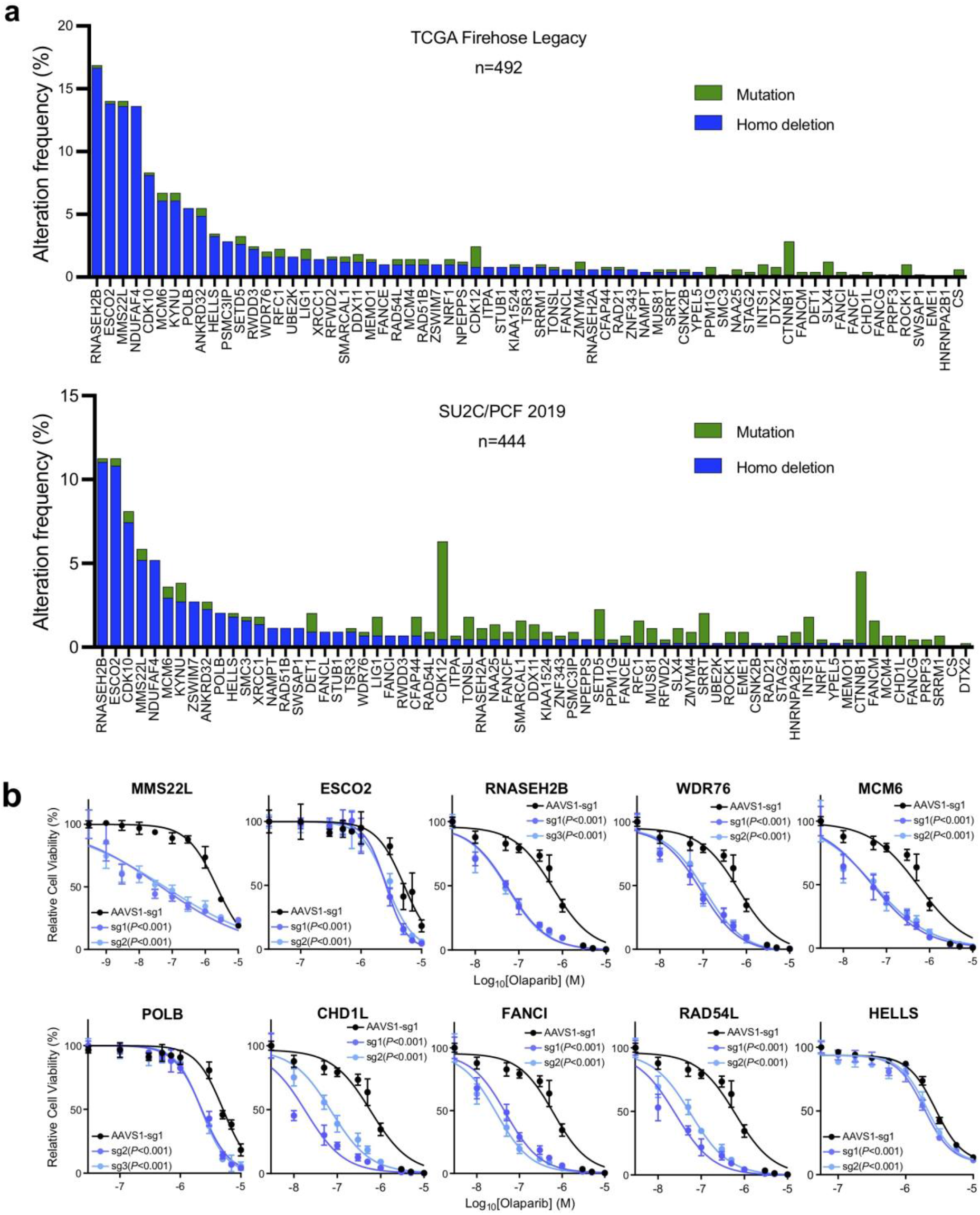
Validation of negatively selected genes with frequent genomic alterations. **a,** The frequency of mutations and homozygous (Homo) deletions in 67 common negatively selected genes from the TCGA Firehose Legacy cohort (Upper panel, n = 492) and the SU2C/PCF cohort (Lower panel, n = 444). **b,** Dose response curves after treatment with olaparib in C4-2B cells with knockout (KO) of the indicated genes. The *p*-values were determined by comparing two gene-specific sgRNAs to a control AAVS1 sgRNA using two-way ANOVA. Error bars represent SD, n = 6.

### Loss of MMS22L increases PARPi response due to impaired HRR function in PCa cells

Since MMS22L always forms a complex with TONSL for replication-associated DNA damage repair, TONSL was unsurprisingly identified as one of the top hits as well (Fig. 1I; Supplementary Table 2). Nevertheless, the frequency of MMS22L homozygous deletions (14% in primary and 5% in metastatic prostate tumors) makes it an attractive biomarker to predict PARPi response in PCa. To further validate this finding, we deleted MMS22L and TONSL in five PCa cell lines - LNCaP, C4-2B, 22Rv1, PC-3, DU145 and MDAPCa2b (Fig. 3a). We showed that deletion of either MMS22L or TONSL resulted in significantly increased sensitivity to olaparib in LNCaP, C4-2B, MDAPCa2b and 22Rv1 cells, but not in DU145 or PC-3 cells. In a growth competition experiment, we observed that olaparib treatment significantly reduced the fraction of MMS22L-KO cells in contrast to corresponding control cells when they were equally pre-mixed and grown under the same treatment condition (Fig. 3b). Notably, MMS22L-deleted C4-2B cells were also sensitive to other PARPis (rucaparib, talazoparib, and veliparib), but displayed lower sensitivity to carboplatin, a DNA crosslinking agent (Fig. 3c). In addition, we generated MMS22L-KO C4-2B cell clones through single cell sorting and confirmed either complete (sg1 clone #1-4) or partial MMS22L deletion (sg1 clone #5-6; sg2, clone #1-6) using immunoblot (Fig. 3d). All MMS22L-KO clones exhibited high sensitivity to olaparib to a similar extent without a gene dose effect, supporting MMS22L haploinsufficiency for its function. Restoration of MMS22L expression completely abolished the sensitivity of MMS22L-KO cells to olaparib (Fig. 3e), confirming that the phenotype was specifically due to MMS22L loss.

**Fig. 3.**
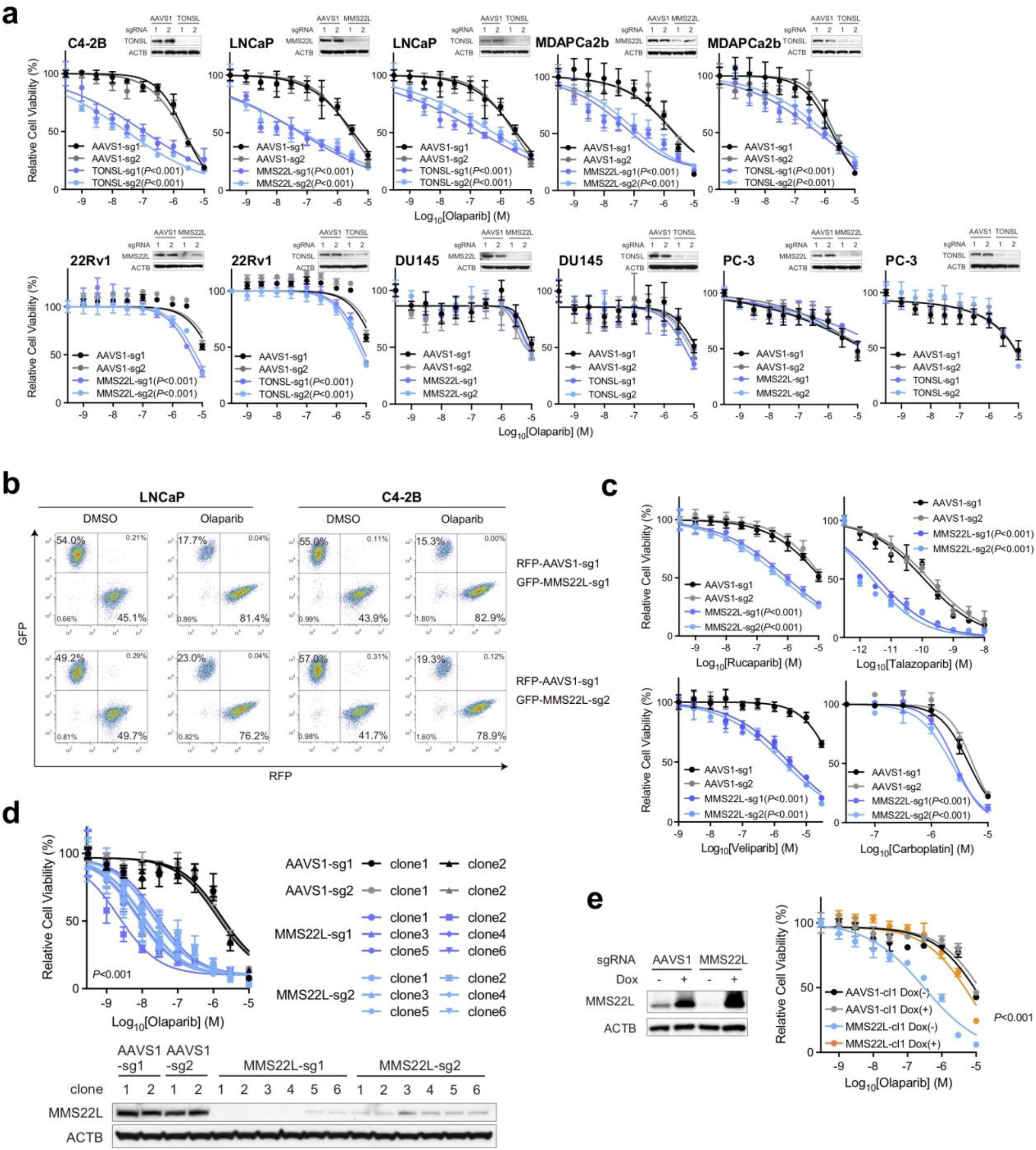
Loss of MMS22L increases PARPi response in PCa cells. **a,** Dose response curves after treatment with olaparib in MMS22L- or TONSL-KO cells versus corresponding AAVS1 control cells of C4-2B, LNCaP, MDAPCa2b, 22Rv1, DU145 and PC-3. Upper right panels in each cell line are immunoblot analysis of MMS22L or TONSL KO efficiency. ACTB (β-actin) is a loading control. The olaparib response in MMS22L-KO C4-2B cells is presented in (**2b**). **b,** Flow cytometry analyses of GFP and RFP positive cells. MMS22L-KO LNCaP or C4-2B cells (with GFP) were co-cultured with corresponding AAVS1 control cells (with RFP) in a 1:1 ratio in the presence of DMSO or olaparib (3 μM). Two MMS22L-KO cell lines (sg1 and sg2) and one control cell line (sg1) were used. Cells were collected and analyzed using flow cytometry 7 days after the treatment. The percentage of each cell population is presented in each panel. **c,** Dose response curves after treatment with rucaparib, talazoparib, veliparib and carboplatin in two MMS22L-KO C4-2B cell lines (sg1 and sg2) versus two AAVS1 control cell lines (sg1 and sg2). **d,** Dose response curves (upper panel) after treatment with olaparib in control and MMS22L-KO C4-2B cell clones. Immunoblot analysis (lower panel) of the MMS22L protein level in AAVS1 control and MMS22L-KO cell clones. **e,** Immunoblot analysis (left panel) of the MMS22L protein level in C4-2B AAVS1 control sg1 clone 1 (cl1) and MMS22L-KO sg1 clone 1 (cl1) stably infected with TET-inducible sgRNA-resistant MMS22L gene, after treatment with or without doxycycline (0.15 μg/ml) for 3 days. Dose response curves (right panel) after olaparib treatment with or without doxycycline (0.15 μg/ml) treatment in the same C4-2B cell clones. The *p*-value was determined by comparison between the MMS22L-KO cell clone with and without doxycycline treatment using two-way ANOVA. Error bars represent SD, n = 6. In (**a**), (**c**) and (**d**), the *p*-values were determined by comparing MMS22L- or TONSL-KO cells to corresponding control cells using two-way ANOVA. Error bars represent SD, n = 6.

Next, we sought to investigate the mechanism by which loss of MMS22L increased PARPi sensitivity. We recently reported that PARPi induces replication fork collapse and DSBs in a trans cell cycle manner and BRCA1/2-deficient cells cannot recruit RAD51 to repair them, leading to cell death^20^. Similarly, we observed increased expression of γ-H2AX and cleaved-PARP in MMS22L-KO C4-2B cells following 72h olaparib treatment (Fig. 4a), reflecting PARPi-induced DNA DSBs and apoptosis. Increased γ-H2AX foci were detected by immunofluorescence staining (Fig. 4b). Cell cycle analysis showed a substantial accumulation of cells in G2/M phase after deletion of MMS22L or TONSL (Fig. 4c). Given the function of the MMS22L-TONSL complex in loading RAD51 at DSB sites and promoting HRR after replication fork collapse independent of BRCA2^38^, we postulated that loss of MMS22L might impair HRR that occurred predominantly in the late S and G2 phases of the cell cycle, leading to overwhelming DSBs and mitotic catastrophe. Indeed, using immunofluorescence staining, we found significantly reduced RAD51 foci in MMS22L-KO C4-2B cells after olaparib treatment for 24 hours in comparison with control cells (Fig. 4d). RAD51 loading was enhanced after MMS22L expression was restored. From these observations, we conclude that MMS22L loss confers high sensitivity to PARPis due to compromised RAD51 recruitment to PARPi-induced DSBs, causing homologous recombination deficiency (HRD).

**Fig. 4.**
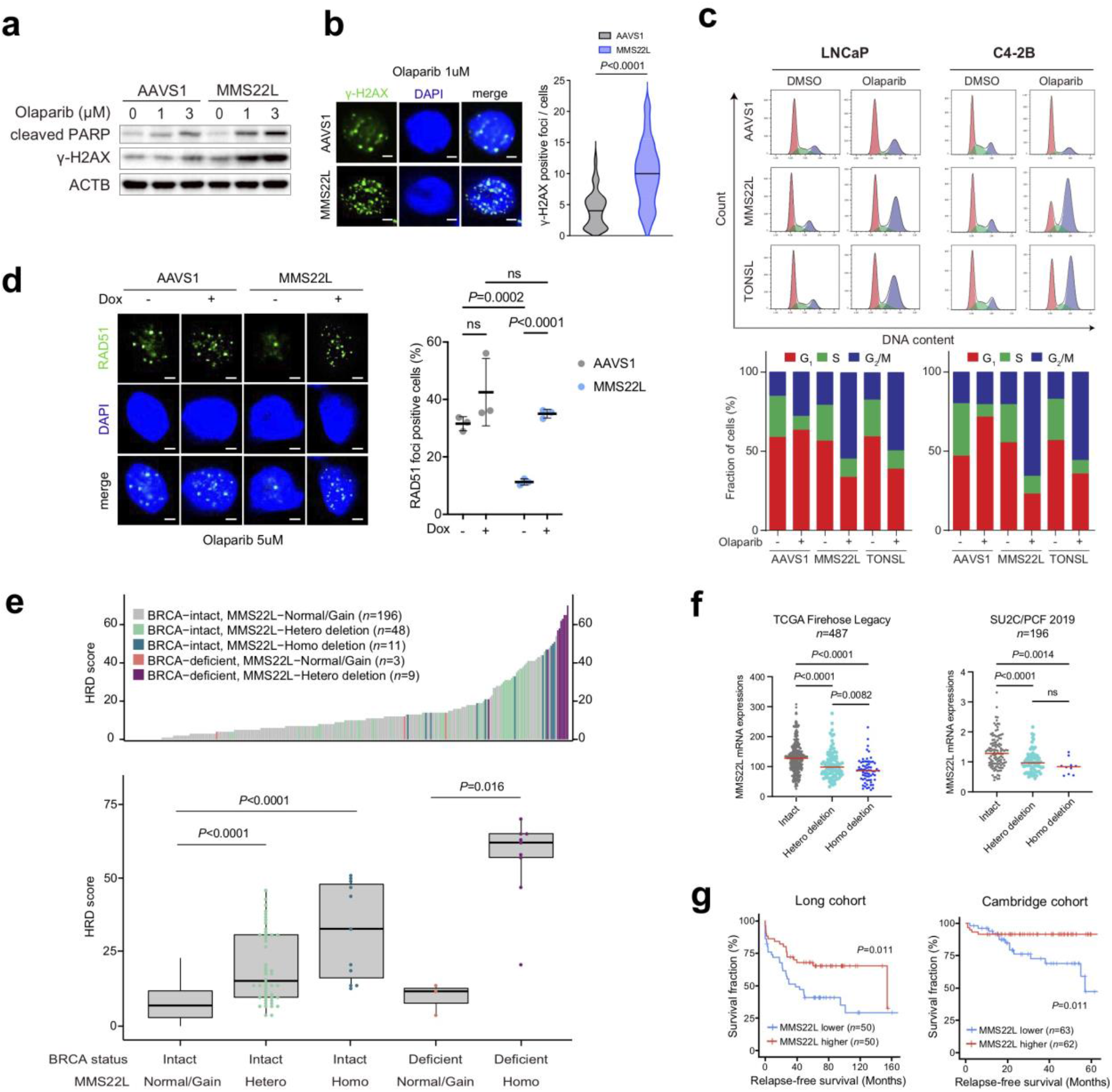
Loss of MMS22L impairs HRR function in PCa cells. **a,** Immunoblot analysis of cleaved PARP and γ-H2AX in AAVS1 control and MMS22L-KO C4-2B cells after olaparib treatment for 72 hours with the indicated concentrations. **b,** Representative images and quantification of γ-H2AX foci in AAVS1 control and MMS22L-KO C4-2B cells after olaparib treatment (1 μM) for 24 hours. More than 100 cells were analyzed per condition. Solid lines inside the violin indicate the median. The *p*-value was determined using unpaired *t*-test. Scale bar = 5 μm. **c,** Cell cycle analysis (upper panel) in AAVS1 control, MMS22L-KO and TONSL-KO LNCaP and C4-2B cells after treatment with DMSO or olaparib (5 μM) for 72 hours. Percentage of cells (lower panel) in each phase of the cell cycle in these cells. **d,** Representative images and quantification of RAD51 foci in AAVS1 control and MMS22L-KO C4-2B cells stably infected with TET-inducible MMS22L gene after olaparib (5 μM) treatment in the presence or absence of doxycycline (0.15 μg/ml) for 24 h. Dots indicate each replicate with more than 100 cells analyzed. The *p*-values were determined using unpaired *t*-test. Error bars represent SD, n = 3. ns = not significant. Scale bar = 5 μm. **e,** Ranked HRD scores (Upper panel) in PCa tumors with BRCA and/or MMS22L genomic alterations as indicated. Comparison of HRD scores (lower panel) between patients groups classified based on BRCA and MMS22L status. HRD score is calculated using whole-genome sequencing datasets as previously described^44^. The *p*-values were determined using unpaired *t*-test. **f,** The mRNA levels of MMS22L in PCa tumors with Intact, heterozygous (Hetero) deletion and homozygous (Homo) deletion of MMS22L in the TCGA cohort (Firehose Legacy; n = 487) and the SU2C/PCF cohort (n = 196). The *P*-values were determined using unpaired *t*-test. **g,** Kaplan-Meier survival curves in the Long PCa cohort^97^ and the Cambridge PCa cohort^98^ based on the MMS22L mRNA expression level (lower versus higher). A log-rank test was carried out to examine the survival difference.

Tumors with HRD have been found to display specific patterns of genomic alterations, which can be quantified using a HRD score based on genome-wide measuring three patterns of genomic instability (loss of heterozygosity, telomeric allelic imbalance, and large-scale state transitions)^41^. A high HRD score (≥42) has been shown to be predictive of clinical benefit with PARPi or platinum therapy in breast and ovarian cancer patients^42,43^. Using whole-genome sequencing data, we previously showed that a subset of PCa patients with a high HRD score did not harbor germline or somatic mutations in BRCA1/2 and other HRR genes^44^. Further analysis revealed that tumors with MMS22L heterozygous or homozygous deletion displayed significantly higher HRD scores (Fig. 4e), although much higher scores were observed in tumors with homozygous deletion. Tumors with both BRCA and MMS22L deficiency had the highest HRD score. The MMS22L loss-mediated genomic instability was further confirmed using HRDetect (Supplementary Fig. 6), a model that quantitatively aggregates six HRD-associated signatures^45^. These results may partially explain previously uncharacterized cause of HRD in PCa. In addition, using publicly available PCa clinical data, we found MMS22L deletion is correlated with decreased MMS22L transcript levels (Fig. 4f). Tumors with heterozygous or homozygous deletion had significantly lower MMS22L mRNA levels compared to those with wild-type MMS22L. Patients with lower MMS22L expression had a shorter survival (Fig. 4g), suggesting tumor progression driven by HRD-mediated genomic instability.

### The response to PARP inhibition after MMS22L loss is p53-dependent

Next, we asked why deletion of MMS22L did not increase PARPi response in PC-3 and DU145 cells (Fig. 3a). CRISPR screens revealed that TP53 was one of the top resistance genes in LNCaP, C4-2B, and 22Rv1 cells (Fig. 1i; Supplementary Table 4). This was validated by deletion of TP53 in these cell lines (Fig. 5a). LNCaP and C4-2B cells possess wild-type TP53, while 22Rv1 cells harbor a mono-allelic TP53 mutation and have a relatively low p53 protein level (Supplementary Table 1 and Supplementary Fig. 7). In contrast, PC-3 cells have TP53 truncating mutations and do not express p53 (p53-null), and DU145 cells harbor dominant negative TP53 mutations^46^. Notably, PCa cell lines with TP53 mutations are generally less sensitive to olaparib compared to cell lines with wild-type TP53 (Supplementary Fig. 1). Furthermore, RNA-sequencing (RNA-seq) analysis revealed 271 commonly upregulated genes in LNCaP and C4-2B cells after olaparib treatment, in which the p53 pathway is significantly enriched (Fig. 5b,c; Supplementary Table 6), supporting the role of p53 activation in modulating response to PARP inhibition. We reasoned that TP53 status might impact PARPi response of MMS22L-deleted cells. Using an RNA interference approach, we showed that knockdown of MMS22L expression significantly increased PARPi sensitivity in TP53 wild-type C4-2B cells, whereas TP53-KO cells remained insensitive (Fig. 5d). Using a Dox-inducible system, we reintroduced the TP53 gene into p53-null PC-3 cells. Restoration of p53 expression resensitized MMS22L-KO PC-3 cells to olaparib *in vitro* and *in vivo* (Fig. 5e, f). We observed that the level of γ-H2AX protein expression and foci were increased after restoration of p53 expression in MMS22L-KO PC-3 cells, indicating more DNA DSBs (Fig. 5g,h). Accordingly, cell apoptosis was increased as determined by cleaved PARP levels. These results suggest that MMS22L deletion increases PARPi sensitivity only when p53 remains functional. While 6% of primary and 15.5% of metastatic tumors contain concurrent genomic alterations of MMS22L and TP53, approximately 27% of primary and 20.5% of metastatic prostate tumors harbor wild-type TP53 and MMS22L deletion when heterozygous deletion is counted (Fig. 5i). To further identify MMS22L deletion in clinical samples, we utilized DNAscope assay, a new chromogenic DNA *in situ* hybridization technique, on a tissue microarray (TMA) with 146 primary PCa tissue cores. We detected 4.1% homozygous deletion and 26% heterozygous deletion of MMS22L in primary PCa patients (Fig. 5j; Supplementary Fig. 8). Together, our results suggest that loss of MMS22L occurs in a considerable fraction of PCa patients, who may therefore benefit from PARP inhibition.

**Fig. 5.**
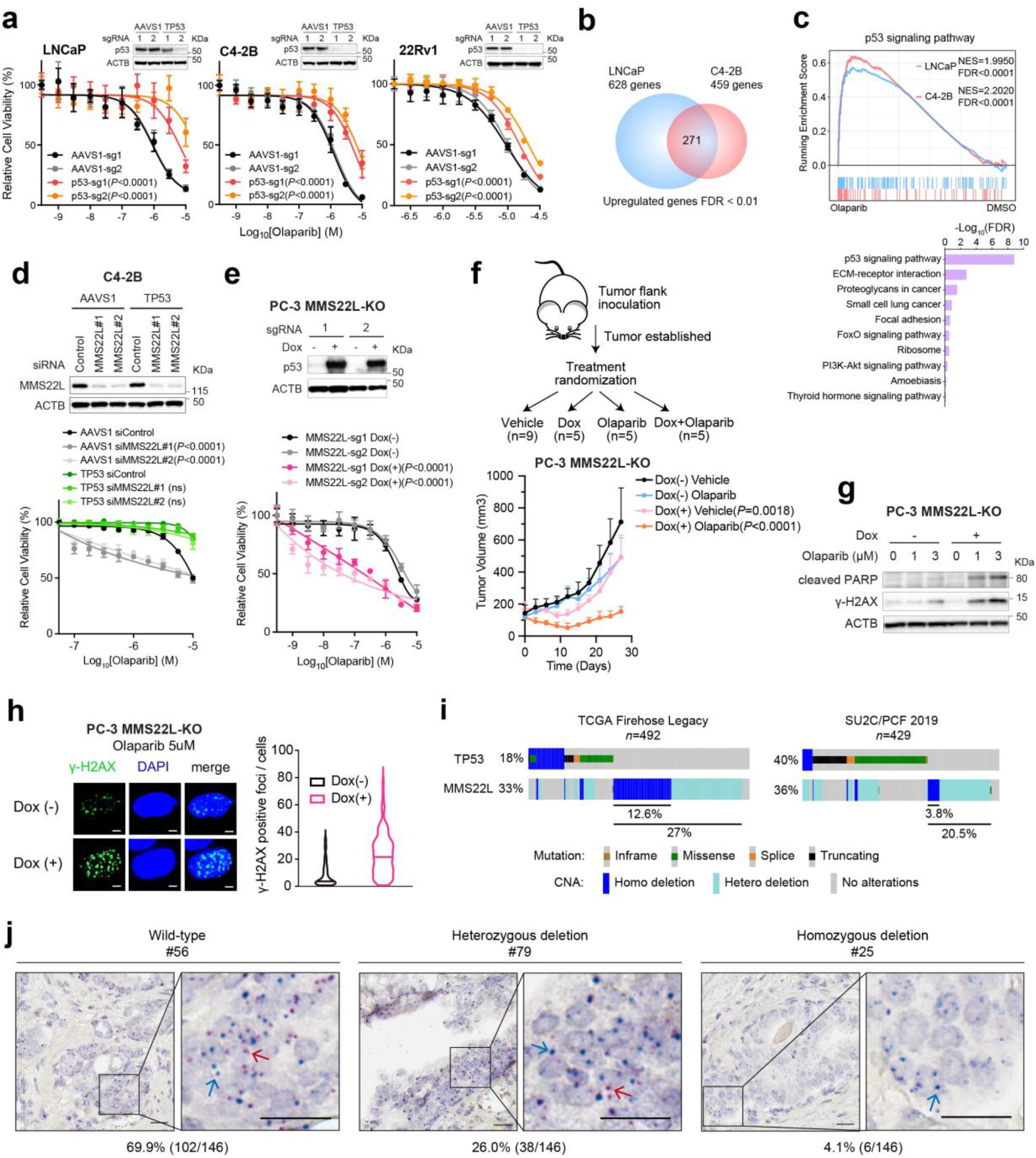
TP53 status impacts PARPi response in MMS22L-depleted PCa cells. **a,** Dose response curves after treatment with olaparib in LNCaP, C4-2B, and 22Rv1 cells after TP53 deletion. The *p*-values were determined by comparing two TP53-KO cell lines (sg1 and sg2) to two corresponding AAVS1 control cell lines (sg1 and sg2) using two-way ANOVA. Error bars represent SD, n = 6. **b,** Venn diagram of up-regulated genes in LNCaP and C4-2B cells after olaparib treatment. **c,** GSEA (upper panel) of RNA-seq data showing enrichment of p53 signaling pathway in up-regulated genes in LNCaP and C4-2B after olaparib treatment. Top KEGG pathways (lower panel) enriched in these up-regulated genes. Normalized enrichment score (NES) and false discover rate (FDR) in each cell line are shown. **d,** Immunoblot analysis (upper panel) of the MMS22L protein level after siRNA knockdown. Dose response curves (lower panel) after olaparib treatment in TP53-KO and AAVS1 control C4-2B cells with or without MMS22L siRNA knockdown. Two different MMS22L siRNAs (#1 and #2) and one control siRNA were transfected into TP53-KO and control C4-2B cells for 2 days followed by olaparib treatment for 7 days. The *p*-values were determined by comparing siMMS22L to siControl using two-way ANOVA. Error bars represent SD, n = 6. **e,** Immunoblot analysis (upper panel) of p53 in AAVS1 control and MMS22L-KO PC-3 cells stably infected with TET-inducible TP53 gene in the presence or absence of doxycycline (0.15 μg/ml) for 3 days. Dose response curves (lower panel) after olaparib treatment for 7 days in the presence or absence of doxycycline (0.15 μg/ml) in these cells. The *p*-values were determined by comparing between cells treated with and without doxycycline using two-way ANOVA. Error bars represent SD, n = 6. **f,** Schematic of the procedure (upper panel) in the xenograft model using MMS22L-KO PC-3 cells containing TET-inducible TP53 gene. Tumor growth (lower panel) in ICR-SCID mice after treatment with olaparib (50 mg/kg) or vehicle with or without doxycycline induction for 4 weeks. The *p*-values were determined by comparing between treatment with and without doxycycline using two-way ANOVA. Error bars represent SD, n = 5. **g,** Immunoblot analysis of cleaved PARP and γ-H2AX in MMS22L-KO PC-3 cells containing TET-inducible TP53 gene after the treatment as indicated for 3 days. **h,** Representative images and quantification of γ-H2AX foci in AAVS1 control and MMS22L-KO PC-3 cells containing TET-inducible TP53 gene after olaparib (5 μM) treatment for 24 h in the presence or absence of doxycycline (0.15 μg/ml). More than 100 cells were analyzed per condition. Solid lines inside the violin indicate the median. The *p*-value was determined by unpaired *t*-test. Scale bar = 5 μm. **i,** Frequency of genomic alterations in the TP53 and MMS22L genes in the TCGA cohort (left panel; Firehose Legacy, n = 492) and the SU2C/PCF cohort (right panel; n = 429). **j,** Representative images (40X) of MMS22L wildtype (WT), heterozygous (Hetero) deletion, and homozygous (Homo) deletion determined by DNAscope using a tissue microarray (TMA) containing 146 primary prostate tumor samples. Red signals (red arrow) indicate a probe targeting the MMS22L gene on chromosome 6q. Blue signals (blue arrow) indicate a chromosome enumeration control probe targeting the centromeric region of chromosome 6p (CEP6p). The number of red and blue dots were counted and the ratio of red/blue was calculated. WT, red/blue > 0.5; Hetero deletion, red/blue between 0.1-0.5; Homo deletion, red/blue < 0.1. Scale bar = 20 μm.

### Validation of positively selected genes related to PARPi resistance

Next, we set out to investigate PARPi resistance. We found that the top positively selected genes include those previously identified as conferring PARPi resistance in other contexts, including PARP1, PARP2, PARG, ADPRHL2 (also known as ARH3), and TP53BP1 (Fig. 1i)^3,47–50^. Although not altered as frequently as TP53 in PCa, downregulation of these genes could influence PARPi response. PPP2R2A was previously included in the HRR gene list used to select patients for the PROfound trial (NCT02987543). However, the FDA subsequently removed PPP2R2A from the approved HRR gene panel because outcomes appeared worse for patients with PPP2R2A alterations with olaparib compared to the control^6^. In accordance with the clinical result, we identified PPP2R2A as one of the common hits from positive selection (Fig. 1i; Supplementary Table 4). PPP2R2A is located on chromosome 8p and frequently deleted in PCa (Supplementary Fig. 9). Indeed, chromosome 8p is the most frequent lost chromosomal arm of all tumors (36%) in the TCGA cohort^35,51,52^, suggesting a potentially common intrinsic resistance mechanism in tumors with 8p loss.

Surprisingly, we identified Checkpoint kinase 2 (CHEK2) as one of the top resistance genes in LNCaP, C4-2B and 22Rv1 cells. This is an unexpected result since CHEK2 is one of the previously reported BRCAness genes^53^ and an FDA-approved biomarker for olaparib in the treatment of mCRPC patients. CHEK2 has been used as a biomarker of PARPi sensitivity in several PARPi clinical trials^7–10,54,55^ because of its known function in promoting HRR through phosphorylation of BRCA1^56,57^. We deleted CHEK2 in five PCa cell lines and observed significantly increased resistance in LNCaP, C4-2B, 22Rv1 cells, but not in TP53-mutant PC-3 and DU145 cells using cell viability and colony formation assays (Fig. 6a,b). We further generated CHEK2-KO C4-2B single cell clones, which all displayed resistance to olaparib (Fig. 6c). The resistance was also observed with other PARPis (rucaparib, talazoparib and veliparib) and carboplatin using C4-2B and 22Rv1 cell models (Fig. 6d). We conclude that CHEK2 loss is associated with PARPi resistance rather than sensitivity in PCa with functional TP53.

**Fig. 6.**
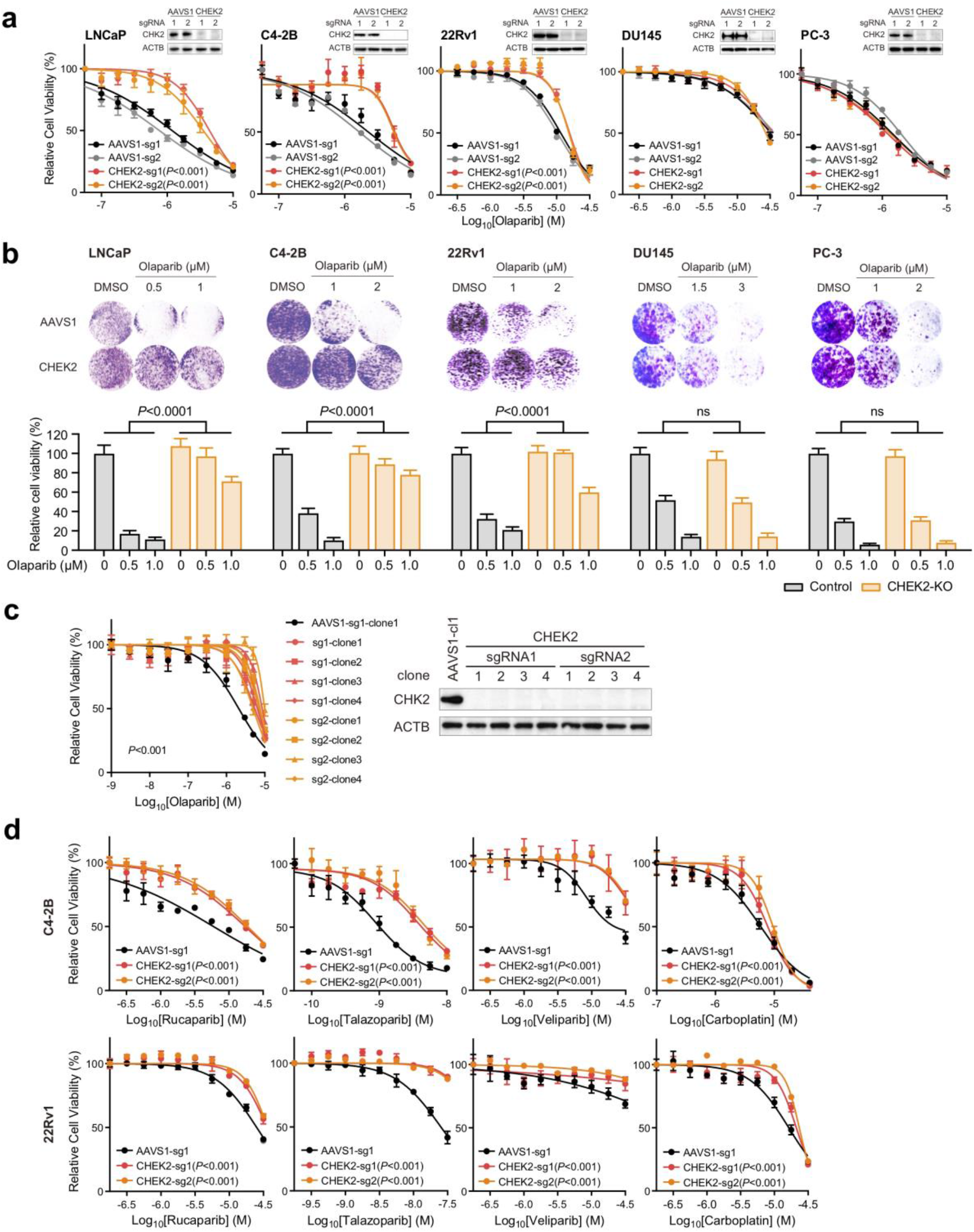
Loss of CHEK2 renders PCa cells resistant to PARP inhibition. **a,** Dose response curves after treatment with olaparib in two CHEK2-KO (sg1 and sg2) and two AAVS1 control (sg1 and sg2) LNCaP, C4-2B, 22Rv1, DU145, and PC-3 cell lines. Upper right panel in each cell line is Immunoblot analysis of the CHEK2 protein level in CHEK2-KO versus control cells. **b,** Representative colony growth images (upper panel) and quantification (lower panel) after treatment with olaparib in CHEK2-KO and AAVS1 control PCa cell lines as indicated. The *p*-values were determined using one-way ANOVA. Error bars represent SD, n = 3. **c,** Dose response curves (left panel) after treatment with olaparib in CHEK2-KO and AAVS1 control C4-2B cell clones. Immunoblot analysis (right panel) of the CHEK2 protein level in CHEK2-KO and control cell clones. **d,** Dose response curves after treatment with rucaparib, talazoparib, veliparib and carboplatin in CHEK2-KO and AAVS1 control C4-2B and 22Rv1 cells. In (**a**), (**c**) and (**d**), the *p*-values were determined by comparing CHEK2-KO to AAVS1 control cells using two-way ANOVA. Error bars represent SD, n = 6.

### Loss of CHEK2/TP53 enhances HRR function through E2F7-controlled BRCA2 expression

CHEK2 regulates multiple proteins beyond BRCA1 in response to DNA damage. CHEK2 phosphorylates p53 on serine 20 and stabilizes p53, leading to cell cycle arrest in G1 phase^58,59^. We therefore asked whether PARPi resistance caused by CHEK2 loss is mediated through p53 inactivation. In line with activation of p53 pathway after PARP inhibition (Fig. 5c), we found that olaparib treatment increased phosphorylated CHEK2 and p53 protein expression levels in CHEK2-intact C4-2B and 22Rv1 cells, but not in CHEK2-KO cells (Fig. 7a). The extent of DNA DSBs was also greater in CHEK2-intact cells compared to CHEK2-KO cells following olaparib treatment as measured by γ-H2AX protein expression and foci (Fig. 7a,b). Accordingly, there was increased olaparib-induced apoptosis in CHEK2-intact cells as measured by cleaved-PARP levels (Fig. 7a). Deletion of TP53 had no effect on olaparib-induced CHK2 phosphorylation but reduced DNA DSBs (γ-H2AX foci) and cell apoptosis (cleaved-PARP) as similarly observed in CHEK2-KO cells (Fig. 7c).

**Fig. 7.**
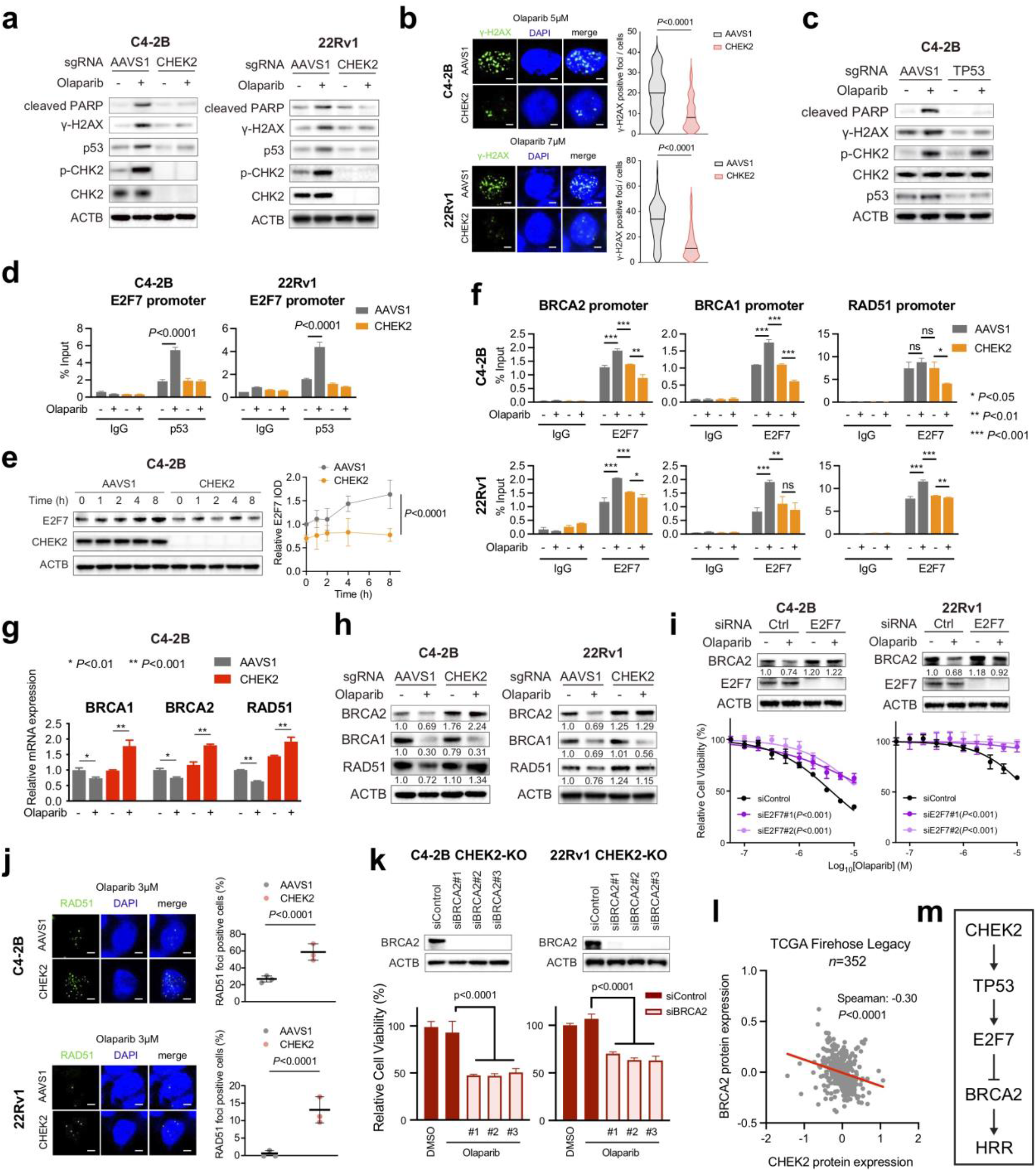
Loss of CHEK2/TP53 enhances HRR function through E2F7-controlled BRCA2 expression. **a,** Immunoblot analysis of the indicated proteins in CHEK2-KO and AAVS1 control C4-2B and 22Rv1 cells after treatment with and without olaparib (3 μM for C4-2B and 5 μM for 22Rv1) for 72 h. **b,** Representative images (left panel) and quantification (right panel) of γ-H2AX foci in CHEK2-KO and AAVS1 control C4-2B and 22Rv1 cells after treatment with olaparib (5 μM for C4-2B and 7 μM for 22Rv1) for 24 h. More than 100 cells were analyzed per condition. Solid lines inside the violin indicate the median. The *p*-values were determined using unpaired *t*-test. Scale bar = 5 μm. **c,** Immunoblot analysis of the indicated proteins in TP53-KO and AAVS1 control C4-2B cells after treatment with and without olaparib (3 μM) for 72 h. **d,** p53 ChIP-qPCR at the E2F7 promoter in CHEK2-KO and AAVS1 control C4-2B and 22Rv1 cells after treatment with or without olaparib (10 μM) for 24 h. **e,** Immunoblot analysis (left panel) of the E2F7 protein level in CHEK2-KO and AAVS1 control C4-2B cells after treatment with olaparib (10 μM) for the indicated time. Quantitative evaluation of integrated optical density (IOD) (right panel) was performed from three independent experiments. The E2F7 IOD values were normalized by ACTB IOD. The results are shown as mean of each time point. The *p*-values were determined by comparing CHEK2-KO to control cells using two-way ANOVA. Error bars represent SD, n=3. **f**, E2F7 ChIP-qPCR at the promoter regions of BRCA1/2 and RAD51 genes in CHEK2-KO and AAVS1 control C4-2B and 22Rv1 cells after treatment with or without olaparib (10 μM) for 24 h. **g**, The mRNA expression of BRCA1/2 and RAD51 genes determined by RT-qPCR in CHEK2-KO and AAVS1 control C4-2B cells after treatment with and without olaparib (10 μM) for 8 h. **h,** Immunoblot analysis of the indicated proteins in CHEK2-KO and AAVS1 control C4-2B and 22Rv1 cells after treatment with and without olaparib (10 μM) for 72 h. The IOD values of the indicated proteins normalized by ACTB IOD are shown. **i,** Immunoblot analysis (upper panel) of BRCA2 expression in C4-2B and 22Rv1 cells after knockdown of E2F7 through RNA interference with or without olaparib (3 μM) treatment for 72 h. Two E2F7 siRNAs and one negative control (Ctrl) siRNA were used (also see Fig. 16). Dose response curves (lower panel) after treatment with olaparib for 7 days in C4-2B and 22Rv1 cells transfected with siRNAs against E2F7 or control. The *p*-values were determined by comparing siE2F7 to siControl using two-way ANOVA. Error bars represent SD, n = 6. **j,** Representative images and quantification of RAD51 foci in CHEK2-KO and AAVS1 control C4-2B cells after treatment with olaparib (3 μM) for 24 h. Dots indicate each replicate with more than 100 cells analyzed. n = 3. Scale bar = 5 μm. **k,** Cell viability after treatment with olaparib (1 μM) for 7 days in CHEK2-KO C4-2B and 22Rv1 cells transfected with siRNAs against BRCA2 or control. n = 6. Immunoblot analysis of the BRCA2 protein level after siRNA knockdown (upper panel). **l,** The correlation between CHEK2 and BRCA2 protein levels in the TCGA cohort (Firehose Legacy; n = 352). Spearman correlation is calculated. **m,** Schematic model of HRR function regulated by the CHEK2-TP53-E2F7-BRCA2 pathway. In (**d**), (**f**), (**g**) (**j**) and (**k**), the *p*-values were determined using unpaired *t*-test. Error bars represent SD. * *p*<0.05, ** *p*<0.01, *** *p*<0.001, ns= not significant.

Next, we sought to investigate the CHEK2-TP53 downstream target genes that may contribute to PARPi resistance in CHEK2-KO cells. Previous studies have shown that E2F7 is a TP53-reulated gene^60^, and that deletion of E2F7 renders cells resistant to PARP inhibition through upregulation of RAD51 in BRCA2-deficient cells^61^. E2F7 was also one of the top positively selected genes in our CRISPR screens (Fig. 1 i). E2F7 is an atypical member of the E2F transcription factor family, which regulates its target genes, including HRR genes, through transcription repression rather than activation. We reasoned that CHEK2 loss might derepress HRR gene expression via the TP53-E2F7 axis. To investigate TP53-mediated regulation of E2F7 expression, we analyzed publicly available p53 ChIP-seq data in multiple cell lines and found highly enriched p53 binding immediately upstream of the E2F7 transcription start site (TSS) (Supplementary Fig. 10a), indicating a conserved and direct transcriptional regulation mechanism. Using p53 ChIP-qPCR, we observed baseline p53 binding at the E2F7 promoter, which was significantly increased after olaparib treatment in CHEK2-intact C4-2B and 22Rv1 cells (Fig. 7d). However, olaparib-induced p53 binding was not observed after CHEK2 loss. Accordingly, E2F7 expression was increased after olaparib treatment in CHEK2-intact C4-2B cells within a few hours, but not in CHEK2-KO cells (Fig. 7e). Furthermore, the MDM2 inhibitor (MDM2i) nutlin, an agent that blocks MDM2-mediated ubiquitination and promotes p53 stabilization and activity, increased p53 expression and p53 binding at the E2F7 promoter as well (Supplementary Fig. 10b,c), supporting p53-regulated E2F7 expression. Indeed, MDM2i treatment overcame olaparib resistance as a result of CHEK2 deletion in line with the role of TP53-E2F7 axis in PARPi resistance (Supplementary Fig. 10d).

To investigate E2F7-controlled HRR gene expression, we analyzed publicly available E2F7 ChIP-seq data in multiple cell lines and observed strong E2F7 binding immediately upstream of the TSSs of BRCA1/2 and RAD51 genes (Supplementary Fig. 10e). Using E2F7 ChIP-qPCR, we demonstrated strong E2F7 binding at these genomic regions in both CHEK2-intact and CHEK2-KO C4-2B and 22Rv1 cells (Fig. 7f). Importantly, E2F7 binding was significantly increased in CHEK2-intact control cells after olaparib treatment but remained unchanged or significantly decreased in CHEK2-KO cells. In agreement with the E2F7 ChIP results, we found that mRNA levels of BRCA1/2 and RAD51 were significantly decreased after olaparib treatment in CHEK2-intact C4-2B cells but increased in CHEK2-KO cells (Fig. 7g), indicating E2F7-mediated transcriptional suppression on HRR genes. It should be noted that no significant cell cycle alterations were observed after CHEK2 or TP53 deletion in C4-2B cells (Supplementary Fig. 11a). Therefore, upregulation of BRCA1/2 and RAD51 gene expression is largely due to transcriptional regulation rather than cell cycle alteration, although the expression of HRR genes is cell cycle-dependent. The protein levels of BRCA1/2 and RAD51 were decreased as well after olaparib treatment in CHEK2-inatct cells (Fig. 7h). However, the protein expression changes were not all consistent with the mRNA expression levels in CHEK2-KO cells. While the BRCA2 and RAD51 protein levels were increased or remained at a high level following olaparib treatment, the BRCA1 protein level was decreased. This is likely because BRCA1 is directly regulated by CHEK2 through posttranslational modification^56^. On the other hand, the reduction of BRCA1 protein expression was not observed in TP53-KO cells since p53 is a downstream effector of CHEK2 (Supplementary Fig. 11b). These results suggest that the CHEK2-TP53-E2F7 axis is activated after olaparib treatment, leading to suppression of HRR gene expression, and that this axis is disrupted when either CHEK2 or TP53 is lost. Specifically, the protein level of the BRCA2 gene was significantly upregulated after CHEK2 or TP53 loss. In addition, we found that olaparib-induced BRCA2 suppression was rescued after knockdown of E2F7, rendering cell resistant to olaparib (Fig. 7i; Supplementary Fig. 11c).

To determine whether HRR capacity was enhanced after CHEK2 loss, we performed RAD51 foci formation assay and found significantly increased olaparib-induced RAD51 foci in CHEK2-KO C4-2B and 22Rv1 cells (Fig. 7j). These results suggest that the PARPi resistance arising from CHEK2 loss is, at least in part, dependent on enhanced HRR function. To further determine whether increased BRCA2 expression is responsible for PARPi resistance after CHEK2 loss, we knocked down BRCA2 in CHEK2-KO C4-2B and 22Rv1 cells and found that these cells were re-sensitized to olaparib (Fig. 7k). Similarly, knockdown of BRCA2 in TP53-KO cells increased olaparib sensitivity (Supplementary Fig. 11d). In agreement with these results, we found that there was a negative correlation between CHEK2 and BRCA2 protein expression in the TCGA cohort (Fig. 7l). Together, our results support the notion that the CHEK2-TP53-E2F7-BRCA2 pathway is one of the primary resistance mechanisms for PARP inhibition after CHEK2 loss (Fig. 7m).

### ATR inhibition overcomes PARPi resistance in CHEK2-deficient PCa cells

Finally, we sought to explore therapeutic strategies to overcome PARPi resistance after CHEK2 loss. Emerging evidence has shown that PARPi resistance is often accompanied by increased ATR activity, which coordinates cell cycle checkpoint response through phosphorylation of CHK1 and allows cells to survive PARPi-induced replication stress^62^. We found that ATR activity was elevated in CHEK2-KO cells after olaparib treatment, as evidenced by increased CHK1 phosphorylation in a dose-dependent manner (Supplementary Fig. 12). We treated CHEK2-KO cells with olaparib in combination with M6620, a clinically used ATR inhibitor (ATRi)^63^. M6620 was remarkably more effective when combined with olaparib in cell viability assays (Fig. 8a). The combination therapy profoundly suppressed cell colony formation of CHEK2-KO cells in a synergistic manner, as demonstrated by the HSA and Bliss synergy scores^64,65^ (Fig. 8b). Using *in vivo* xenograft models, we observed that CHEK2-KO C4-2B and 22Rv1 tumors did not respond to olaparib or M6620 as a single agent. However, the combination treatment abolished the tumor growth (Fig. 8c and Supplementary 13). We did not observe any weight loss in each group, indicating the combination therapy was well-tolerated. Mechanistically, M6620 inhibited CHK1 phosphorylation as expected (Fig. 8d). The combination treatment significantly inhibited BRCA1/2 and RAD51 expression in CHEK2-KO C4-2B and 22Rv1 cells, leading to more DNA damage and cell apoptosis as determined by γ-H2AX and cleaved-PARP expression. Functionally, olaparib-induced RAD51 formation was completely abolished in CHEK2-KO cells when they were treated with M6620 in combination with olaparib (Fig. 8e).

**Fig. 8.**
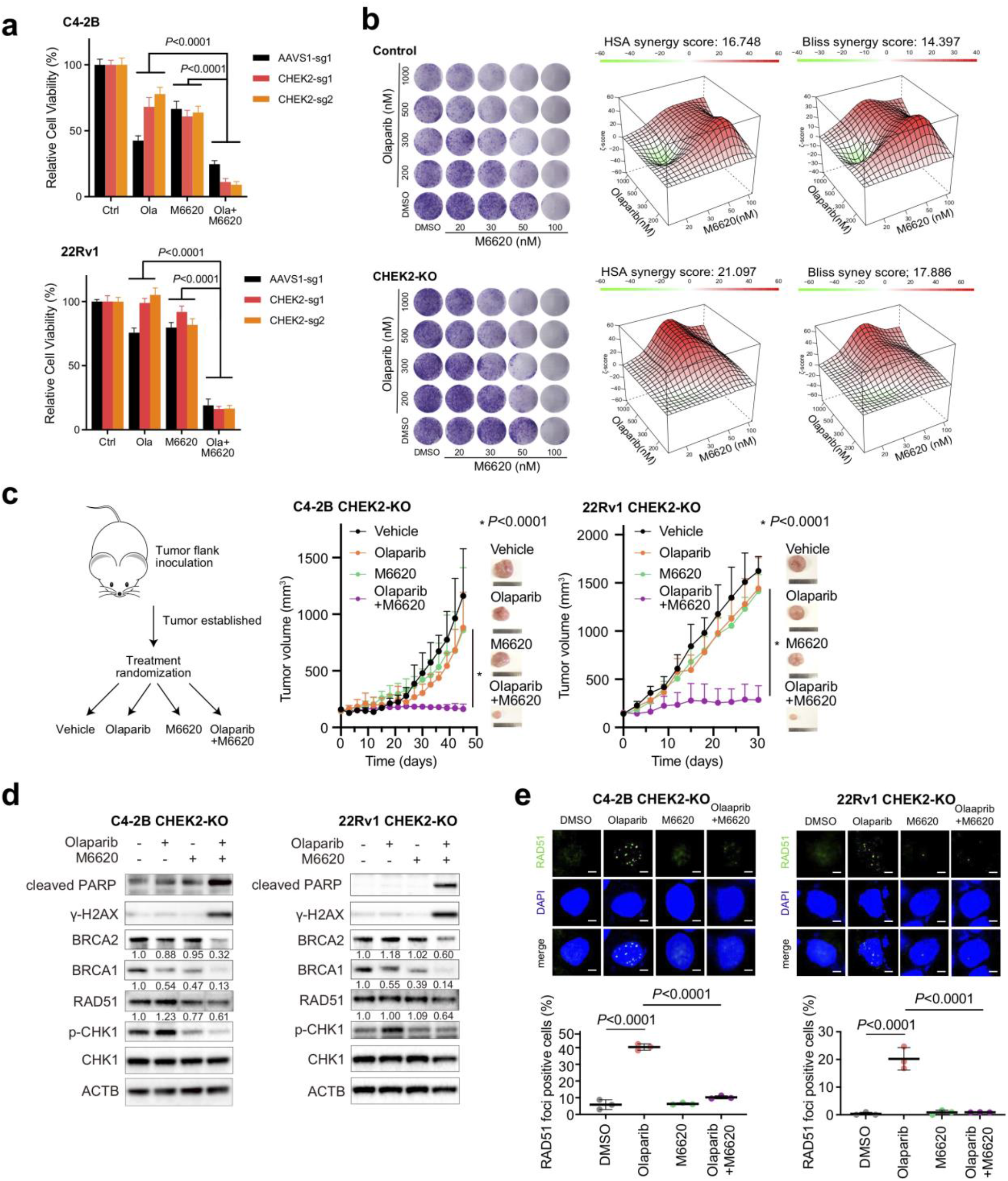
ATR inhibition overcomes PARPi resistance caused by CHEK2 loss in PCa cells. **a,** Cell viability in CHEK2-KO and AAVS1 control C4-2B and 22Rv1 cells after treatment with DMSO, olaparib (3 μM), M6620 (100 nM) or combination of olaparib and M6620. The *p*-values were determined using one-way ANOVA. Error bars represent SD, n = 6. **b,** Representative colony growth images (left panel) after combination treatment of olaparib and M6620 in CHEK2-KO and AAVS1 control C4-2B cells. 3D synergy maps (right panel) of HSA and Bliss scores between olaparib and M6620. **c,** Schematic of the procedure (left panel) in xenograft mouse models using CHEK2-KO C4-2B and 22Rv1 cells. After tumors developed to a volume of around 150 mm^3^, mice were randomized into 4 treatment groups as vehicle, olaparib (50 mg/kg), M6620 (25 mg/kg) and olaparib plus M6620. The single agent was administered 5 days/week, while the combination treatment was given 5 days/week for olaparib plus 4 days/week for M6620. Tumor growth of CHEK2-KO C4-2B (middle panel) and 22Rv1 (right panel) cells in ICR-SCID mice treated as indicated are shown with representative images. * p < 0.0001 was determined using two-way ANOVA. Error bars represent SD, n = 4. **d,** Immunoblot analyses of the indicated proteins in CHEK2-KO C4-2B and 22Rv1 cells after treatment as indicated (olaparib, 3 μM; M6620, 100 nM) for 72 h. The IOD values of the indicated proteins normalized by ACTB IOD were shown. **e,** Representative images (upper panel) and quantification (lower panel) of RAD51 foci in CHEK2-KO C4-2B and 22Rv1 cells after treatment as indicated (olaparib, 10 μM; M6620, 200 nM) for 24 h. Dots indicate each replicate with more than 100 cells analyzed. The *p*-values were determined using unpaired *t*-test. Error bars represent SD, n = 3. Scale bar = 5 μm.

## DISCUSSION

In this study, we provided a systematic view of the genetic determinants and mechanisms underlying the sensitivity and resistance of PCa to PARP inhibition beyond BRCA1/2 alterations. We identified genes, such as RAD51B, RAD54L, FANCL, that have already been used as biomarkers for PARP inhibition in the FDA-approved genetic test for mCRPC patients, as well as previously unknown genes. These new genes could serve as predictive biomarkers for PARP inhibition if mutated or deleted in PCa or therapeutic targets through pharmacologic inhibition in combination with PARPis. One such novel hit is MMS22L, a gene located in a genomic region frequently deleted in PCa, loss of which significantly increases the response to PARPis. Mechanistically, MMS22L forms a complex with TONSL, which accumulates at distressed replication forks and is required for the HRR of replication fork-associated DSBs through promoting RAD51 loading^38,39,66^. When MMS22L is deleted, cells fail to load RAD51 to PARPi-induced collapsed replication forks, leading to accumulation of DSBs, cell cycle arrest at G2/M phase, and apoptotic cell death. The HRR function of MMS22L-TONSL seems specific to the recovery of collapsed replication forks and does not act through BRCA2^38^. Interestingly, MMS22L deletions appear to be an early event in prostate tumorigenesis since they are more frequently detected in primary tumors. If the use of PARPis in localized PCa (e.g., in the neoadjuvant/adjuvant setting) is found to confer clinical benefit as has been suggested in breast cancer with BRCA mutations^67^, detection of MMS22L deletion would identify a much larger population of patients who may benefit. In addition, our data suggest that cells with partial loss of MMS22L (i.e., monoallelic loss) are equally sensitive to PARP inhibition compared to complete loss (i.e., biallelic loss) likely due to insufficient MMS22L-TONSL complex formation. This is further supported by the evidence from clinical sample analyses, showing significantly lower mRNA levels and higher HRD scores in both homozygous and heterozygous MMS22L deletion tumors. Identification of MMS22L as a new BRCAness gene explains a significant part of patients that have high HRD scores but without HRR gene alterations per the olaparib label. Given a considerable number of PCa patients with MMS22L deletion detected by next generation sequencing- or DNA *in situ* hybridization-based assays, this genomic alteration may be a valuable biomarker for PARP inhibition despite the fact that the response is p53-dependent.

Previous studies of PARPi resistance have been focusing on acquired resistance mechanisms in BRCA1/2-deficient cancers, showing that residual or restored HRR activity is the most commonly observed resistance mechanism. This can be achieved by secondary or reversion mutations of BRCA1, BRCA2 and RAD51 isoforms^68–70^, loss of BRCA1 promoter methylation^71^, increased expression of hypomorphic isoforms of BRCA1^72,73^, loss of TP53BP1 and resection-associated factors RIF1, REV7, and Shieldin^49,74–76^. In contrast, our work uncovers an intrinsic resistance mechanism in BRCA1/2-sufficient PCa cells, involving two frequently mutated tumor suppressor genes TP53 and CHEK2. The role of TP53 and its upstream activator CHEK2 in a set of tightly regulated cell cycle checkpoints and DDR events has been extensively studied. In response to DNA DSBs, cells activate CHEK2 that phosphorylates and stabilizes p53, leading to cell cycle arrest to allow for DNA repair or inducing apoptosis after genotoxic damage. However, the PARPi resistance caused by loss of CHEK2 or TP53 cannot be fully explained by impaired p53-mediated apoptosis. Indeed, PARPi can trigger apoptotic cell death in TP53-deficient PCa cells^77^. High-grade serous ovarian cancer (HGSOC) patients with defects in HRR are highly sensitive to PARP inhibition despite the fact that TP53 mutations are detected in 96% HGSOC tumors^78^. Here, we propose an as yet uncharacterized pathway that underlies the resistance of PCa cells to PARP inhibition. We show that PARP inhibition activates the CHEK2-TP53-E2F7 pathway that suppresses the expression of HRR genes and potentiates the cytotoxicity of PARPis. Loss of CHEK2 or TP53, however, markedly reduces the expression of their downstream target E2F7, leading to increased HRR gene (largely BRCA2) expression due to lack of E2F7-mediated transcriptional suppression. Upregulation of BRCA2 enhances HRR capacity sufficient for the repair of PARPi-induced DSBs and cell survival. Interestingly, BRCA1 is another direct target of the CHEK2 kinase. The fact that loss of CHEK2 leads to PARPi resistance instead of sensitivity is unexpected. We show that loss of CHEK2 may compromise BRCA1 regulation; after all, the BRCA1 protein expression is not completely abolished. On the other hand, the CHEK2-TP53-E2F7 pathway has predominantly emerged after PARP inhibition. Our finding is consistent with the results from recent clinical trials which show little benefit in patients with mutations in CHEK2^8,10,79^, providing a rationale to revisit FDA-approved clinically-used biomarkers for the use of olaparib. Our data support the notion that HRR deficiency resulting from alterations in non-BRCA genes is unlikely to be of similar therapeutic relevance in comparison to BRCA2 deficiency, considering that mutations in the TP53 gene occur in more than 50% of all cancers. Conversely, PCa harboring BRCA2 deficiency more likely respond to PARP inhibition regardless of TP53 status. This is consistent with our CRISPR screen results, showing that BRCA2 was identified as one of the top hits in TP53-mutated DU145 cells, while many of the canonical HRR genes (including Fanconi anemia genes) were not negatively selected. Further investigation is needed to determine which genomic alterations in the HRR pathway can or cannot be rescued by upregulation of BRCA2 after loss of CHEK2 or TP53. We recently reported that RB1, another frequently mutated tumor suppressor gene, may also confer resistance to PARP inhibition through E2F1-mediated upregulation of HRR genes^36^, indicating the importance and complexity of E2F transcriptional network through activation and repression in the context of DNA repair. The finding of the TP53/E2F7- and RB1/E2F1-mediated resistance mechanisms is clinically relevant since TP53 and RB1 are concurrently altered in 39% of mCRPC tumors with adenocarcinoma histology and 74% of mCRPC tumors with neuroendocrine features^80,81^. Current clinical use of PARPis is guided by mutations of a single gene and overlooks common concurrent genomic alterations that may have an antagonistic effect.

Finally, we propose a therapeutic approach with combined PARP and ATR inhibition to overcome PARPi resistance. Mechanistically, in order to survive PARP inhibition, cancer cells rely more on ATR checkpoint to slow down cell cycle and reduce replication stress for repairing DSBs. ATR inhibition increases replication origin firing and DNA synthesis despite the presence of PARPi-induced replication fork gaps, leading to progressive accumulation of DSBs^20^. Moreover, ATR inhibition can disrupt RAD51 loading to stalled replication forks and DSBs in PARPi-resistant BRCA1-deficient cells^82^. Our results further demonstrate that blocking ATR activity inhibits HRR gene expression and RAD51 foci formation, leading to a high number of stalled replication forks collapsing into toxic, irreparable DSBs. Indeed, the combination of PARPi with ATRi was synergistic in many PARPi-resistant models regardless of their resistance mechanisms^62^. Simultaneous suppression of ATR and PARP may be a preferred strategy to overcome PARPi resistance.

## METHODS

### Cell Lines and Materials

Prostate cancer cell lines LNCaP, C4-2B, 22Rv1, PC-3, DU145, and MDA PCa 2b were obtained from American Type Culture collection (ATCC). Cells were cultured following the provider’s recommendations. The identity of the cell lines was confirmed based on high-resolution small tandem repeats (STR) profiling at Dana-Farber Cancer Institute (DFCI) Molecular Diagnostics Core Laboratory and exome-sequencing analyses performed by LC Sciences. Cells were regularly tested for mycoplasma. All reagents (including antibodies, small molecule inhibitors, oligonucleotides, and plasmids) used in this study are listed in Supplementary Table 7.

### Cellular Viability Assay

PCa cells were seeded in 96-well plates with 1000-3000 cells/well (n = 6) and treated as indicated. Cellular viability assays were performed using alamarBlue (ThermoFisher Scientific) 5-10 days after treatments and repeated 2–3 times.

### Clonogenic Assay

PCa cells were seeded in 12-well plates at low density to avoid contact between clones. Subsequently, cells were treated as indicated and allowed to grow for 10-14 days. Colonies were fixed with paraformaldehyde (4%) for 20 minutes and stained with crystal violet (0.1%) for 15 minutes. Colony images were quantified using ImageJ software (National Institutes of Health).

### Cell Growth Competition Assay

MMS22L-KO and AAVS1 control cells were infected with lentiviruses generated using LV-GFP (Addgene #25999) or LV-RFP (Addgene #26001) plasmid, respectively. MMS22L-KO LNCaP and C4-2B cells stably express GFP, whereas control cells stably express RFP. GFP or RFP positive cells were collected by the BD FACSAria cell sorter (BD Biosciences) and mixed in a 1:1 ratio. The mixed cell populations were incubated for 7 days with 3 uM of olaparib or DMSO treatment, followed by collection and calculation of GFP and RFP positive cells.

### Drug Synergy analysis

Cells were seeded in 12-well plates and treated with vehicle and 4 doses of olaparib (200, 300, 500 and 1000 nM) and M6620 (20, 30, 50 and 100 nM) in a matrix format to include 25 different dose combinations. Colonies were quantified on day 14 using imageJ. Drug synergy scores were calculated based on the HSA and Bliss model using the SynergyFinder2.0^64^.

### RNA Interference

E2F7 and BRCA2 siRNAs and negative control siRNA were purchased from Sigma-Aldrich. Cells were seeded onto 6-well or 96-well plates for 24 hours and followed by siRNA (10 nM) transfection using Lipofectamine RNAiMAX (Invitrogen) according to the manufacturer’s protocol. The siRNA sequences are listed in Supplementary Table 7.

### Cell Cycle Analysis

Cells were plated in 6-cm dishes with or without olaparib treatment for 72 hours. Collected cells were labelled using Click-iT^®^ EdU Flow Cytometry Assay Kit (Invitrogen) according to the manufacturer’s protocol and followed by flow cytometry using BD LSR Fortessa (BD Biosciences).

### Chromatin Immunoprecipitation Quantitative PCR (ChIP-qPCR)

ChIP experiments were performed as previously described^83,84^. Briefly, PCa cells were grown in regular media and treated with olaparib (10 μM) for 24 hours prior to ChIP. Cells were cross-linked by formaldehyde (1%) at room temperature (RT) for 10 minutes. After washing with icecold PBS, cells were collected and lysed. The soluble chromatin was purified and fragmented by sonication. Immunoprecipitation (IP) was performed using normal IgG or p53 or E2F7 antibody (2 μg/IP). ChIP DNA was extracted and analyzed by qPCR using iTaq Universal SYBR Green Supermix (Bio-Rad). The primer sequences are listed in Supplementary Table 7.

### Quantitative RT-PCR (RT-qPCR)

Total RNA was extracted using RNeasy Mini Kit (Qiagen) according to the manufacturer’s protocol. RT-qPCR was performed as previously described^85^. The primer sequences are listed in Supplementary Table 7.

### RNA Sequencing (RNA-seq)

LNCaP and C4-2B cells were treated with olaparib (3 μM) or DMSO in biological duplicates. Total RNAs were extracted using RNeasy Mini Kit (Qiagen). RNA-Seq library preparation and next generation sequencing by Illumina HiSeq were conducted using GENEWIZ services.

### *In vivo* Xenograft studies

All mice were maintained in compliance with the guidelines approved by the Institutional Animal Care and Use Committee (IACUC) at the Brigham and Women’s Hospital. For xenograft studies, male ICR/SCID mice and male NCG mice (*NOD-Prkdc^em26Cd52^Il2rg^em26Cd22^/NjuCrl*) at 4-5 weeks of age were purchased from Tconic Bioscience and Charles River Laboratories, respectively. For the PC-3 xenograft assay, 2.5 million cells were prepared in 50 μl PBS and mixed with 50 μl Matrigel (Corning Matrigel Matrix High Concentration #354262) to form a total of 100 μl cell suspension and followed by the subcutaneous inoculation in male ICR/SCID mice. Tumor volume was calculated using the modified ellipsoid formula: length x width^2^/2. After tumors developed to a volume of approximately150 mm^3^, mice were randomized into 4 groups (vehicle: n = 9, doxycycline: n = 5, olaparib: n = 5, and doxycycline plus olaparib: n = 5). Mice were treated with vehicle (5% DMSO/10% d-a-tocopherol polyethylene glycol 1000 succinate; oral gavage, OG), doxycycline (10 mg/kg in 10% d-a-tocopherol polyethylene glycol 1000 succinate; intraperitoneal, IP), olaparib (50 mg/kg in 5%DMSO/10% d-a-tocopherol polyethylene glycol 1000 succinate; OG) or doxycycline (IP) plus olaparib (OG). Treatment was administered every day for doxycycline and 5 days a week for vehicle and olaparib.

For the C4-2B and 22Rv1 xenograft assays, 5 million cells of C4-2B or 2.5 million cells of 22Rv1 were prepared as above. After tumors developed to a volume of approximately150 mm^3^, mice were randomized into 4 groups (vehicle, olaparib, M6620, and combination treatment) with 4 mice in each group. Mice were treated with vehicle, olaparib, M6620 (25 mg/kg in 5%DMSO/10% d-a-tocopherol polyethylene glycol 1000 succinate) or combination treatment (olaparib 50 mg/kg and M6620 25 mg/kg in 5%DMSO/10% d-a-tocopherol polyethylene glycol 1000 succinate) through OG. Treatment was administered 5 days a week for single agent, and 5 days a week of olaparib plus 4 days a week of M6620 for combination treatment. Tumor volume was manually measured every 3 days.

### Immunoblot analysis

Cells were lysed in RIPA buffer supplemented with Halt™ protease and phosphatase inhibitor cocktail (ThermoFisher Scientific). The protein concentration was determined using Pierce BCA Protein Assay Kit (ThermoFisher Scientific). Proteins were resolved in SDS-polyacrylamide gels (4–12%) and transferred to PVDF membranes using a Tris-glycine buffer system. Membranes were blocked with 5% non-fat milk in 0.1% Tween20 in TBS (TBS-T) for 1 hour at room temperature followed by incubation with primary antibodies and subsequently corresponding secondary antibodies in 5% milk TBS-T. The membranes were developed with Immobilon substrate (EMD Millipore).

### PAR immunoblotting

The assay was performed according to the protocol as previously described^48^. Briefly, cells were grown on 10 cm dishes for 24 hours to achieve 90% confluency. Cells were treated with olaparib and/or PDDX-001 in the presence or absence of 0.01 % MMS for a short period of time as indicated. For total protein extraction, cells were lysed in RIPA buffer supplemented with protease and phosphatase inhibitor cocktail and 1 μM PARG inhibitor ADP-HPD (Merck). For chromatin fraction, cells were lysed and each fraction was collected using Subcellular Protein Fractionation Kit (ThermoFisher Scientific) according to the manufacturer’s protocol. Immunoblotting was carried out as described in above.

### Immunofluorescence Analyses

Cells were grown in a poly-L lysine-coated 4 well Millicell EZ slides (EMD Millipore) for 48 hours and pre-extracted with 0.2% Triton-X on ice 90 seconds before fixation with 4% paraformaldehyde in PBS for 20 minutes at room temperature (RT) and then permeabilized by incubation with ice-cold methanol. After permeabilization, cells were blocked with 5% BSA in PBS for 1 hour at RT, followed by incubation with 5% BSA in TBS-T containing primary antibodies at a ratio of 1:200 overnight at 4°C. Cells were washed and incubated with secondary fluorescent antibodies for 1 hour at RT. Alexa Fluor 488 anti-rabbit and Alexa Fluor 488 antimouse antibodies were purchased from Invitrogen. After washing, the nuclear content was stained with Mounting Medium with DAPI (Abcam) overnight at 4°C. Images were obtained with fluorescence microscope BX53 (Olympus). Images were quantitatively assessed using ImageJ software (NIH, Bethesda, MD) with ‘Find maxima’ function. More than 100 cells were analyzed per condition in each experiment in a blinded manner.

### Generating Knockout Cell Lines Using CRISPR/Cas9 Gene Editing

CRISPR guides targeting each gene were cloned into lentiGuide-Puro vector (#52963; Addgene). The sgRNA sequences are listed in the Supplementary Table 7. The lentiCas9-Blast vector that expresses Cas9 was obtained from Addgene (#52962). Lentiviruses were generated using packaging vectors pMD2.G (#12259; Addgene) and psPAX2 (#12260; Addgene) with Lipofectamine™3000 transfection reagents (#L3000015; Invitrogen) in 293FT cells. PCa cells were infected with lentiviruses expressing Cas9 and selected with Blasticidin (10 μg/ml) to establish stable Cas9-expressing cell lines. Polybrene was added at a final concentration of 8ug/ml to increase infecting efficiency. To generate KO cells, PCa cells were infected with lentiviruses containing specific sgRNA and selected with puromycin (3 μg/ml). KO efficacy was determined by immunoblot analyses. Single cell clones of MMS22L-KO and CHEK2-KO were generated using BD FACSAria cell sorter (BD Biosciences) and followed by expansion.

### Generation of TET-inducible Exogenous MMS22L- or P53-expressing Cells

MMS22L-KO and control C4-2B clone cells were infected with viruses containing TET-inducible MMS22L gene. The MMS22L cDNA was created to be resistant to MMS22L sgRNA1 by the introduction of silent mutations in the crispr RNA (crRNA) recognition sequence (change CTTGGCAGGAATATAGCACAA to **CTAGGTAGA**AATATAGCACAA).

After neomycin (500 mg/ml) selection, doxycycline (0.15 μg/ml) treatment induces the MMS22L expression confirmed by immunoblot. In addition, MMS22L-KO PC-3 cells were infected with viruses containing TET-inducible WT TP53. Neomycin selection and the confirmation of p53 expression were performed as above. The MMS22L and TP53 cDNAs were cloned into the Lenti-TRE3G-ORF-IRES-tRFP-PGK-Tet3G-neo vector, respectively. The plasmids were custom-designed and synthesized at Transomic Technologies, Inc.

### Genome-wide CRISPR-Cas9 Knockout Screen

We used genome-wide CRISPR-Cas9 KO H1 and H2 libraries obtained from Drs. Myles Brown and X. Shirley Liu’s laboratories^15^, consisting of over 180,000 sgRNAs (10sgRNAs per gene). For the CRISPR-Cas9 KO screens, 200 million PCa cells were infected with the lentiviral CRISPR-Cas9 KO H1 and H2 libraries at a low multiplicity of infection (~0.3) to to maximize the number of cells that have only one sgRNA integration. After 5 days of puromycin selection, the surviving cells were expanded. Following preparation of 60 million cells for each condition to achieve a representation of at least 300 cells per sgRNA, cells were divided into day 0 control cells and cells cultured for 28 days (nine passages) treated with DMSO or olaparib before genomic DNA extraction and library preparation. We used olaparib at the concentration of 5 μM for LNCaP and C4-2B cells and 10 μM for 22Rv1 and DU145 cells. These concentrations were close to the half maximal inhibitory concentration (IC50) for each cell line and allowed us to identify both negatively (depleted) and positively (enriched) selected sgRNAs corresponding to gene knockouts that increase and decrease olaparib response respectively. PCR was performed using genomic DNA to construct the sequencing libraries. Each library was sequenced at 30-40 million reads to achieve ~300 x average coverage over the CRISPR library. The library from day 0 sample of each screen served as controls to identify positively or negatively selected genes.

### CRISPR-Cas9 KO Screen Data Analysis

CRISPR-Cas9 KO screen data was analyzed using the MAGeCK and MAGeCK-VISPR algorithms^86,87^. MAGeCK calculated the read counts for each sgRNA. MAGeCK-VISPR calculated the β-score for each gene by using AAVS1 gene as control. Comparison of the differential β-score between olaparib treatment and DMSO treatment was performed using MAGeCKFlute^17,18^. We ranked genes by differential β-score and robustly estimated σ, which is the standard deviation of the differential β-score by a “quantile matching” approach. A cut-off value was set to correspond 99 % of the data falling within 2σ, which defined genes with lower β-scores than minus σ and higher β-scores than σ as negatively and positively selected genes, respectively. Identified genes were further analyzed using STRING protein interaction analysis^21^. GO analysis was performed using DAVID bioinformatics resources (https://david.ncifcrf.gov).

### DNAscope Assay Using Tissue Microarray (TMA)

The TMA was constructed using primary prostate tumors retrieved from the Vancouver Prostate Centre Tissue Bank as previously reported as approved by the institutional Review Board^88^. A total of 146 tissue cores from 73 patients who had undergone radical prostatectomy were included in this study. The DNAscope assay (Advanced Cell Diagnostics, Newark, CA) is a chromogenic DNA in situ hybridization assay using two sets of target specific probes. Two custom-designed probes were synthesized by Advanced Cell Diagnostics. DS-Hs-MMS22L-C1 is the probe targeting 29194-49180 of hg38 DNA range = chr 6:97142161-97283437. DS-Hs-CEP6p-C2 is the control probe targeting 20891-35730 of hd38 DNA range = chr 6: 57224489-57312704. The assay was performed on the TMA using DNAscope™ HD Duplex Reagent Kit (Advanced Cell Diagnostics) according to the manufacturer’s protocol. Briefly, the tissue section was pretreated to allow access to target DNAs and followed by hybridization with two sets of probes. Two independent signal amplification systems were used to detect both target DNAs. Probes were hybridized to a cascade of signal amplification molecules, culminating in binding of enzyme-labeled probes. Two chromogenic substrates were used and probe targeted regions were visualized in red for the MMS22L locus and blue for the control region (i.e., the centromeric region of chromosome 6p). An increase in the number of red dots relative to blue dots indicates a copy number gain or amplification, while a decrease in the number of red dots or no red dots indicates a copy number loss or deletion. The images were evaluated and scored both digitally by the software (Aperio ImageScope, Leica Biosystems) and visually by a pathologist (L. Fazli, Vancouver Prostate Centre). The MMS22L wild-type was defined as the ratio of the total number of red dots divided by the total number of blue dots (red/blue) greater than 0.5. The heterozygous deletion was defined as the ratio of red/blue is between 0.1-0.5, while the homozygous deletion was defined as the ratio of red/blue is below 0.1.

### RNA Sequencing Analysis

Raw RNA-seq reads are aligned to the human genome version hg38 by using STAR aligner^89^. Gene counts are quantified by using HT-seq^90^ with REFSEQ annotation. Differentially expressed genes are identified by using DESeq2^91^ with cutoff of FDR < 0.01, and ranked by the statistics. The GSEA software was used for determining the KEGG pathways^92^.

### HRD and HRDetect Scores

HRD and HRdetect scores were determined as previously described^44^. A total of 267 PCa cases with whole genome sequencing data were analyzed using the scarHRD R package and HRDetect algorithm^45,93^.

### Statistical Analysis

Statistical analyses were performed using the unpaired 2-tailed Student’s *t* test or 2-way ANOVA with a post hoc Tukey’s honest significant difference (HSD) test when comparing at least three conditions using the Prism software (GraphPad). *P*-values of less than 0.05 were considered as statistically significant.

### Public Data Analysis

Clinical data sets were analyzed using the cBio Portal for Cancer Genomics (cBioPortal; www.cbioportal.org) and PCaDB^94^ (http://bioinfo.jialab-ucr.org/PCaDB/). Integrated Genome Viewer (https://www.broadinstitute.org/igv/) was used for visualization of ChIP-seq data analysis. ChIP-seq data were obtained from ChIP-Atlas^95^ (http://chip-atlas.org).

## Data Availability

RNA-seq data are available under GEO accession number GSE189186.

## Author contribution

T.Tsujino and T.Takai performed the majority of experiments and analyses. T.Tsujino and T.Takai, and K.H undertook the CRISPR screens. T.Tsujino and L.J. wrote the manuscript and prepared figures. F.G. performed animal studies. T.Tsutsumi, X.B., C.M, C.F., B.G., and A.S. performed cell studies and provided technical assistance. Z.Sztupinszki performed HRD analysis. N.X., L.F. and X.D. performed DNA scope assay. H.A. and A.S.K. provided administrative support. T.Tsujino, K.H, A.D.C., K.W.M., Z. Szallasi, L.Z., A.S.K., L.J. analyzed the data and revised manuscript. L.J. conceptualized, designed the study, and acquired funding for the present study.

## Notes

### Competing Interest Statement

The authors have declared no competing interest.

### Summary of Updates

We corrected the errors in Figure 5j and Supplementary Figure 8. The total case number has been changed from 111 to 146. Accordingly, the sample numbers have changed in the figures. No changes were made in the text.

## References

1 Robinson, D. et al. Integrative Clinical Genomics of Advanced Prostate Cancer. Cell 162, 454, doi:10.1016/j.cell.2015.06.053 (2015).

2 Armenia, J. et al. The long tail of oncogenic drivers in prostate cancer. Nat Genet 50, 645–651, doi:10.1038/s41588-018-0078-z (2018).

3 Murai, J. et al. Trapping of PARP1 and PARP2 by Clinical PARP Inhibitors. Cancer Res 72, 5588–5599, doi:10.1158/0008-5472.CAN-12-2753 (2012).

4 Farmer, H. et al. Targeting the DNA repair defect in BRCA mutant cells as a therapeutic strategy. Nature 434, 917–921, doi:10.1038/nature03445 (2005).

5 Bryant, H. E. et al. Specific killing of BRCA2-deficient tumours with inhibitors of poly(ADP-ribose) polymerase. Nature 434, 913–917, doi:10.1038/nature03443 (2005).

6 Hussain, M. et al. Survival with Olaparib in Metastatic Castration-Resistant Prostate Cancer. N Engl J Med 383, 2345–2357, doi:10.1056/NEJMoa2022485 (2020).

7 de Bono, J. et al. Olaparib for Metastatic Castration-Resistant Prostate Cancer. N Engl J Med 382, 2091–2102, doi:10.1056/NEJMoa1911440 (2020).

8 de Bono, J. S. et al. Talazoparib monotherapy in metastatic castration-resistant prostate cancer with DNA repair alterations (TALAPRO-1): an open-label, phase 2 trial. Lancet Oncol 22, 1250–1264, doi:10.1016/S1470-2045(21)00376-4 (2021).

9 Abida, W. et al. Rucaparib in Men With Metastatic Castration-Resistant Prostate Cancer Harboring a BRCA1 or BRCA2 Gene Alteration. J Clin Oncol 38, 3763–3772, doi:10.1200/JCO.20.01035 (2020).

10 Abida, W. et al. Non-BRCA DNA Damage Repair Gene Alterations and Response to the PARP Inhibitor Rucaparib in Metastatic Castration-Resistant Prostate Cancer: Analysis From the Phase II TRITON2 Study. Clin Cancer Res 26, 2487–2496, doi:10.1158/1078-0432.CCR-20-0394 (2020).

11 Mateo, J. et al. Olaparib in patients with metastatic castration-resistant prostate cancer with DNA repair gene aberrations (TOPARP-B): a multicentre, open-label, randomised, phase 2 trial. Lancet Oncol 21, 162–174, doi:10.1016/S1470-2045(19)30684-9 (2020).

12 Olivieri, M. et al. A Genetic Map of the Response to DNA Damage in Human Cells. Cell 182, 481–496 e421, doi:10.1016/j.cell.2020.05.040 (2020).

13 Zimmermann, M. et al. CRISPR screens identify genomic ribonucleotides as a source of PARP-trapping lesions. Nature 559, 285–289, doi:10.1038/s41586-018-0291-z (2018).

14 Clements, K. E. et al. Identification of regulators of poly-ADP-ribose polymerase inhibitor response through complementary CRISPR knockout and activation screens. Nat Commun 11, 6118, doi:10.1038/s41467-020-19961-w (2020).

15 Jeselsohn, R. et al. Allele-Specific Chromatin Recruitment and Therapeutic Vulnerabilities of ESR1 Activating Mutations. Cancer Cell 33, 173–186 e175, doi:10.1016/j.ccell.2018.01.004 (2018).

16 Xu, H. et al. Sequence determinants of improved CRISPR sgRNA design. Genome Res 25, 1147–1157, doi:10.1101/gr.191452.115 (2015).

17 Wang, B. et al. Integrative analysis of pooled CRISPR genetic screens using MAGeCKFlute. Nat Protoc 14, 756–780, doi:10.1038/s41596-018-0113-7 (2019).

18 Li, Z. et al. CRISPR Screens Identify Essential Cell Growth Mediators in BRAF Inhibitor-resistant Melanoma. Genomics Proteomics Bioinformatics 18, 26–40, doi:10.1016/j.gpb.2020.02.002 (2020).

19 Gupte, R., Liu, Z. & Kraus, W. L. PARPs and ADP-ribosylation: recent advances linking molecular functions to biological outcomes. Genes Dev 31, 101–126, doi:10.1101/gad.291518.116 (2017).

20 Simoneau, A., Xiong, R. & Zou, L. The trans cell cycle effects of PARP inhibitors underlie their selectivity toward BRCA1/2-deficient cells. Genes Dev 35, 1271–1289, doi:10.1101/gad.348479.121 (2021).

21 Szklarczyk, D. et al. The STRING database in 2021: customizable protein-protein networks, and functional characterization of user-uploaded gene/measurement sets. Nucleic Acids Res 49, D605–D612, doi:10.1093/nar/gkaa1074 (2021).

22 Hakem, R., de la Pompa, J. L., Elia, A., Potter, J. & Mak, T. W. Partial rescue of Brca1 (5-6) early embryonic lethality by p53 or p21 null mutation. Nat Genet 16, 298–302, doi:10.1038/ng0797-298 (1997).

23 Ludwig, T., Chapman, D. L., Papaioannou, V. E. & Efstratiadis, A. Targeted mutations of breast cancer susceptibility gene homologs in mice: lethal phenotypes of Brca1, Brca2, Brca1/Brca2, Brca1/p53, and Brca2/p53 nullizygous embryos. Genes Dev 11, 1226–1241, doi:10.1101/gad.11.10.1226 (1997).

24 Adam, S. et al. The CIP2A-TOPBP1 axis safeguards chromosome stability and is a synthetic lethal target for BRCA-mutated cancer. Nat. Cancer 2, 1357–1371, doi: https://doi.org/10.1038/s43018-021-00266-w (2021).

25 Kollarovic, G., Topping, C. E., Shaw, E. P. & Chambers, A. L. The human HELLS chromatin remodelling protein promotes end resection to facilitate homologous recombination and contributes to DSB repair within heterochromatin. Nucleic Acids Res 48, 1872–1885, doi:10.1093/nar/gkz1146 (2020).

26 Spruijt, C. G. et al. Dynamic readers for 5-(hydroxy)methylcytosine and its oxidized derivatives. Cell 152, 1146–1159, doi:10.1016/j.cell.2013.02.004 (2013).

27 Gilmore, J. M. et al. WDR76 Co-Localizes with Heterochromatin Related Proteins and Rapidly Responds to DNA Damage. PLoS One 11, e0155492, doi:10.1371/journal.pone.0155492 (2016).

28 Dayebgadoh, G., Sardiu M. E., Florens, L. & Washburn, M. P. Biochemical Reduction of the Topology of the Diverse WDR76 Protein Interactome. J Proteome Res 18, 3479–3491, doi:10.1021/acs.jproteome.9b00373 (2019).

29 Kluth, M. et al. 13q deletion is linked to an adverse phenotype and poor prognosis in prostate cancer. Genes Chromosomes Cancer 57, 504–512, doi:10.1002/gcc.22645 (2018).

30 Brookman-Amissah, N. et al. Allelic imbalance at 13q14.2 approximately q14.3 in localized prostate cancer is associated with early biochemical relapse. Cancer Genet Cytogenet 179, 118–126, doi:10.1016/j.cancergencyto.2007.08.017 (2007).

31 Dong, J. T., Chen, C., Stultz, B. G., Isaacs, J. T. & Frierson, H. F., Jr. Deletion at 13q21 is associated with aggressive prostate cancers. Cancer Res 60, 3880–3883 (2000).

32 Lu, T. & Hano, H. Deletion at chromosome arms 6q16-22 and 10q22.3-23.1 associated with initiation of prostate cancer. Prostate Cancer Prostatic Dis 11, 357–361, doi:10.1038/pcan.2008.4 (2008).

33 Hyytinen, E. R. et al. Defining the region(s) of deletion at 6q16-q22 in human prostate cancer. Genes Chromosomes Cancer 34, 306–312, doi:10.1002/gcc.10065 (2002).

34 Kluth, M. et al. Deletion lengthening at chromosomes 6q and 16q targets multiple tumor suppressor genes and is associated with an increasingly poor prognosis in prostate cancer. Oncotarget 8, 108923–108935, doi:10.18632/oncotarget.22408 (2017).

35 Stopsack, K. H. et al. Aneuploidy drives lethal progression in prostate cancer. Proc Natl Acad Sci U S A 116, 11390–11395, doi:10.1073/pnas.1902645116 (2019).

36 Miao, C. et al. RB1 loss overrides PARP inhibitor sensitivity driven by RNASEH2B loss in prostate cancer. Sci Adv (in press) (2022).

37 Saredi, G. et al. H4K20me0 marks post-replicative chromatin and recruits the TONSL-MMS22L DNA repair complex. Nature 534, 714–718, doi:10.1038/nature18312 (2016).

38 O’Donnell, L. et al. The MMS22L-TONSL complex mediates recovery from replication stress and homologous recombination. Mol Cell 40, 619–631, doi:10.1016/j.molcel.2010.10.024 (2010).

39 Duro, E. et al. Identification of the MMS22L-TONSL complex that promotes homologous recombination. Mol Cell 40, 632–644, doi:10.1016/j.molcel.2010.10.023 (2010).

40 Piwko, W. et al. The MMS22L-TONSL heterodimer directly promotes RAD51-dependent recombination upon replication stress. EMBO J 35, 2584–2601, doi:10.15252/embj.201593132 (2016).

41 How, J. A. et al. Modification of Homologous Recombination Deficiency Score Threshold and Association with Long-Term Survival in Epithelial Ovarian Cancer. Cancers (Basel) 13, doi:10.3390/cancers13050946 (2021).

42 Telli, M. L. et al. Homologous Recombination Deficiency (HRD) Score Predicts Response to Platinum-Containing Neoadjuvant Chemotherapy in Patients with Triple-Negative Breast Cancer. Clin Cancer Res 22, 3764–3773, doi:10.1158/1078-0432.CCR-15-2477 (2016).

43 Ray-Coquard, I. et al. Olaparib plus Bevacizumab as First-Line Maintenance in Ovarian Cancer. N Engl J Med 381, 2416–2428, doi:10.1056/NEJMoa1911361 (2019).

44 Sztupinszki, Z. et al. Detection of Molecular Signatures of Homologous Recombination Deficiency in Prostate Cancer with or without BRCA1/2 Mutations. Clin Cancer Res 26, 2673–2680, doi:10.1158/1078-0432.CCR-19-2135 (2020).

45 Davies, H. et al. HRDetect is a predictor of BRCA1 and BRCA2 deficiency based on mutational signatures. Nat Med 23, 517–525, doi:10.1038/nm.4292 (2017).

46 Chappell, W. H. et al. p53 expression controls prostate cancer sensitivity to chemotherapy and the MDM2 inhibitor Nutlin-3. Cell Cycle 11, 4579–4588, doi:10.4161/cc.22852 (2012).

47 Makvandi, M. et al. A PET imaging agent for evaluating PARP-1 expression in ovarian cancer. J Clin Invest 128, 2116–2126, doi:10.1172/JCI97992 (2018).

48 Gogola, E. et al. Selective Loss of PARG Restores PARylation and Counteracts PARP Inhibitor-Mediated Synthetic Lethality. Cancer Cell 33, 1078–1093 e1012, doi:10.1016/j.ccell.2018.05.008 (2018).

49 Jaspers, J. E. et al. Loss of 53BP1 causes PARP inhibitor resistance in Brca1-mutated mouse mammary tumors. Cancer Discov 3, 68–81, doi:10.1158/2159-8290.CD-12-0049 (2013).

50 Prokhorova, E. et al. Unrestrained poly-ADP-ribosylation provides insights into chromatin regulation and human disease. Mol Cell, doi:10.1016/j.molcel.2021.04.028 (2021).

51 El Gammal, A. T. et al. Chromosome 8p deletions and 8q gains are associated with tumor progression and poor prognosis in prostate cancer. Clin Cancer Res 16, 56–64, doi:10.1158/1078-0432.CCR-09-1423 (2010).

52 Cai, Y. et al. Loss of Chromosome 8p Governs Tumor Progression and Drug Response by Altering Lipid Metabolism. Cancer Cell 29, 751–766, doi:10.1016/j.ccell.2016.04.003 (2016).

53 Lord, C. J. & Ashworth, A. BRCAness revisited. Nat Rev Cancer 16, 110–120, doi:10.1038/nrc.2015.21 (2016).

54 Stopsack, K. H. Efficacy of PARP Inhibition in Metastatic Castration-resistant Prostate Cancer is Very Different with Non-BRCA DNA Repair Alterations: Reconstructing Prespecified Endpoints for Cohort B from the Phase 3 PROfound Trial of Olaparib. Eur Urol 79, 442–445, doi:10.1016/j.eururo.2020.09.024 (2021).

55 Tung, N. M. et al. TBCRC 048: Phase II Study of Olaparib for Metastatic Breast Cancer and Mutations in Homologous Recombination-Related Genes. J Clin Oncol 38, 4274–4282, doi:10.1200/JCO.20.02151 (2020).

56 Lee, J. S., Collins, K. M., Brown, A. L., Lee, C. H. & Chung, J. H. hCds1-mediated phosphorylation of BRCA1 regulates the DNA damage response. Nature 404, 201–204, doi:10.1038/35004614 (2000).

57 Zhang, J. et al. Chk2 phosphorylation of BRCA1 regulates DNA double-strand break repair. Mol Cell Biol 24, 708–718, doi:10.1128/MCB.24.2.708-718.2004 (2004).

58 Hirao, A. et al. DNA damage-induced activation of p53 by the checkpoint kinase Chk2. Science 287, 1824–1827, doi:10.1126/science.287.5459.1824 (2000).

59 Takai, H. et al. Chk2-deficient mice exhibit radioresistance and defective p53-mediated transcription. EMBO J 21, 5195–5205, doi:10.1093/emboj/cdf506 (2002).

60 Aksoy, O. et al. The atypical E2F family member E2F7 couples the p53 and RB pathways during cellular senescence. Genes Dev 26, 1546–1557, doi:10.1101/gad.196238.112 (2012).

61 Clements, K. E. et al. Loss of E2F7 confers resistance to poly-ADP-ribose polymerase (PARP) inhibitors in BRCA2-deficient cells. Nucleic Acids Res 46, 8898–8907, doi:10.1093/nar/gky657 (2018).

62 Kim, H. et al. Combining PARP with ATR inhibition overcomes PARP inhibitor and platinum resistance in ovarian cancer models. Nat Commun 11, 3726, doi:10.1038/s41467-020-17127-2 (2020).

63 Yap, T. A. et al. Phase I Trial of First-in-Class ATR Inhibitor M6620 (VX-970) as Monotherapy or in Combination With Carboplatin in Patients With Advanced Solid Tumors. J Clin Oncol 38, 3195–3204, doi:10.1200/JCO.19.02404 (2020).

64 Ianevski, A., Giri, A. K. & Aittokallio, T. SynergyFinder 2.0: visual analytics of multi-drug combination synergies. Nucleic Acids Res 48, W488–W493, doi:10.1093/nar/gkaa216 (2020).

65 Ianevski, A., He, L., Aittokallio, T. & Tang, J. SynergyFinder: a web application for analyzing drug combination dose-response matrix data. Bioinformatics 33, 2413–2415, doi:10.1093/bioinformatics/btx162 (2017).

66 Huang, T. H. et al. The Histone Chaperones ASF1 and CAF-1 Promote MMS22L-TONSL-Mediated Rad51 Loading onto ssDNA during Homologous Recombination in Human Cells. Mol Cell 69, 879–892 e875, doi:10.1016/j.molcel.2018.01.031 (2018).

67 Tutt, A. N. J. et al. Adjuvant Olaparib for Patients with BRCA1-or BRCA2-Mutated Breast Cancer. N Engl J Med 384, 2394–2405, doi:10.1056/NEJMoa2105215 (2021).

68 Kondrashova, O. et al. Secondary Somatic Mutations Restoring RAD51C and RAD51D Associated with Acquired Resistance to the PARP Inhibitor Rucaparib in High-Grade Ovarian Carcinoma. Cancer Discov 7, 984–998, doi:10.1158/2159-8290.CD-17-0419 (2017).

69 Lin, K. K. et al. BRCA Reversion Mutations in Circulating Tumor DNA Predict Primary and Acquired Resistance to the PARP Inhibitor Rucaparib in High-Grade Ovarian Carcinoma. Cancer Discov 9, 210–219, doi:10.1158/2159-8290.CD-18-0715 (2019).

70 Pettitt, S. J. et al. Clinical BRCA1/2 Reversion Analysis Identifies Hotspot Mutations and Predicted Neoantigens Associated with Therapy Resistance. Cancer Discov 10, 1475–1488, doi:10.1158/2159-8290.CD-19-1485 (2020).

71 Ter Brugge, P. et al. Mechanisms of Therapy Resistance in Patient-Derived Xenograft Models of BRCA1-Deficient Breast Cancer. J Natl Cancer Inst 108, doi:10.1093/jnci/djw148 (2016).

72 Wang, Y. et al. BRCA1 intronic Alu elements drive gene rearrangements and PARP inhibitor resistance. Nat Commun 10, 5661, doi:10.1038/s41467-019-13530-6 (2019).

73 Wang, Y. et al. The BRCA1-Delta11q Alternative Splice Isoform Bypasses Germline Mutations and Promotes Therapeutic Resistance to PARP Inhibition and Cisplatin. Cancer Res 76, 2778–2790, doi:10.1158/0008-5472.CAN-16-0186 (2016).

74 Xu, G. et al. REV7 counteracts DNA double-strand break resection and affects PARP inhibition. Nature 521, 541–544, doi:10.1038/nature14328 (2015).

75 Noordermeer, S. M. et al. The shieldin complex mediates 53BP1-dependent DNA repair. Nature 560, 117–121, doi:10.1038/s41586-018-0340-7 (2018).

76 Gupta, R. et al. DNA Repair Network Analysis Reveals Shieldin as a Key Regulator of NHEJ and PARP Inhibitor Sensitivity. Cell 173, 972–988 e923, doi:10.1016/j.cell.2018.03.050 (2018).

77 Gonzalez-Billalabeitia, E. et al. Vulnerabilities of PTEN-TP53-deficient prostate cancers to compound PARP-PI3K inhibition. Cancer Discov 4, 896–904, doi:10.1158/2159-8290.CD-13-0230 (2014).

78 Cancer Genome Atlas Research, N. Integrated genomic analyses of ovarian carcinoma. Nature 474, 609–615, doi:10.1038/nature10166 (2011).

79 de Bono, J., Kang, J. & Hussain, M. Olaparib for Metastatic Castration-Resistant Prostate Cancer. Reply. N Engl J Med 383, 891, doi:10.1056/NEJMc2023199 (2020).

80 Mu, P. et al. SOX2 promotes lineage plasticity and antiandrogen resistance in TP53-and RB1-deficient prostate cancer. Science 355, 84–88, doi:10.1126/science.aah4307 (2017).

81 Ku, S. Y. et al. Rb1 and Trp53 cooperate to suppress prostate cancer lineage plasticity, metastasis, and antiandrogen resistance. Science 355, 78–83, doi:10.1126/science.aah4199 (2017).

82 Yazinski, S. A. et al. ATR inhibition disrupts rewired homologous recombination and fork protection pathways in PARP inhibitor-resistant BRCA-deficient cancer cells. Genes Dev 31, 318–332, doi:10.1101/gad.290957.116 (2017).

83 Decker, K. F. et al. Persistent androgen receptor-mediated transcription in castration-resistant prostate cancer under androgen-deprived conditions. Nucleic Acids Res 40, 10765–10779, doi:10.1093/nar/gks888 (2012).

84 Gui, B. et al. Selective targeting of PARP-2 inhibits androgen receptor signaling and prostate cancer growth through disruption of FOXA1 function. Proc Natl Acad Sci U S A 116, 14573–14582, doi:10.1073/pnas.1908547116 (2019).

85 Zheng, D. et al. Secretory leukocyte protease inhibitor is a survival and proliferation factor for castration-resistant prostate cancer. Oncogene 35, 4807–4815, doi:10.1038/onc.2016.13 (2016).

86 Li, W. et al. MAGeCK enables robust identification of essential genes from genome-scale CRISPR/Cas9 knockout screens. Genome Biol 15, 554, doi:10.1186/s13059-014-0554-4 (2014).

87 Li, W. et al. Quality control, modeling, and visualization of CRISPR screens with MAGeCK-VISPR. Genome Biol 16, 281, doi:10.1186/s13059-015-0843-6 (2015).

88 Li, Y. et al. SRRM4 gene expression correlates with neuroendocrine prostate cancer. Prostate 79, 96–104, doi:10.1002/pros.23715 (2019).

89 Dobin, A. et al. STAR: ultrafast universal RNA-seq aligner. Bioinformatics 29, 15–21, doi:10.1093/bioinformatics/bts635 (2013).

90 Anders, S., Pyl, P. T. & Huber, W. HTSeq--a Python framework to work with high-throughput sequencing data. Bioinformatics 31, 166–169, doi:10.1093/bioinformatics/btu638 (2015).

91 Love, M. I., Huber, W. & Anders, S. Moderated estimation of fold change and dispersion for RNA-seq data with DESeq2. Genome Biol 15, 550, doi:10.1186/s13059-014-0550-8 (2014).

92 Subramanian, A. et al. Gene set enrichment analysis: a knowledge-based approach for interpreting genome-wide expression profiles. Proc Natl Acad Sci U S A 102, 15545–15550, doi:10.1073/pnas.0506580102 (2005).

93 Sztupinszki, Z. et al. Migrating the SNP array-based homologous recombination deficiency measures to next generation sequencing data of breast cancer. NPJ Breast Cancer 4, 16, doi:10.1038/s41523-018-0066-6 (2018).

94 Li, R. & Jia, Z., doi:10.1101/2021.06.29.449134 (2021).

95 Oki, S. et al. ChIP-Atlas: a data-mining suite powered by full integration of public ChIP-seq data. EMBO Rep 19, doi:10.15252/embr.201846255 (2018).

96 Conway, J. R., Lex, A. & Gehlenborg, N. UpSetR: an R package for the visualization of intersecting sets and their properties. Bioinformatics 33, 2938–2940, doi:10.1093/bioinformatics/btx364 (2017).

97 Long, Q. et al. Global transcriptome analysis of formalin-fixed prostate cancer specimens identifies biomarkers of disease recurrence. Cancer Res 74, 3228–3237, doi:10.1158/0008-5472.CAN-13-2699 (2014).

98 Ross-Adams, H. et al. Integration of copy number and transcriptomics provides risk stratification in prostate cancer: A discovery and validation cohort study. EBioMedicine 2, 1133–1144, doi:10.1016/j.ebiom.2015.07.017 (2015).

