## Supplementary Figures for "CRISPR screens reveal genetic determinants of PARP inhibitor sensitivity and resistance in prostate cancer"

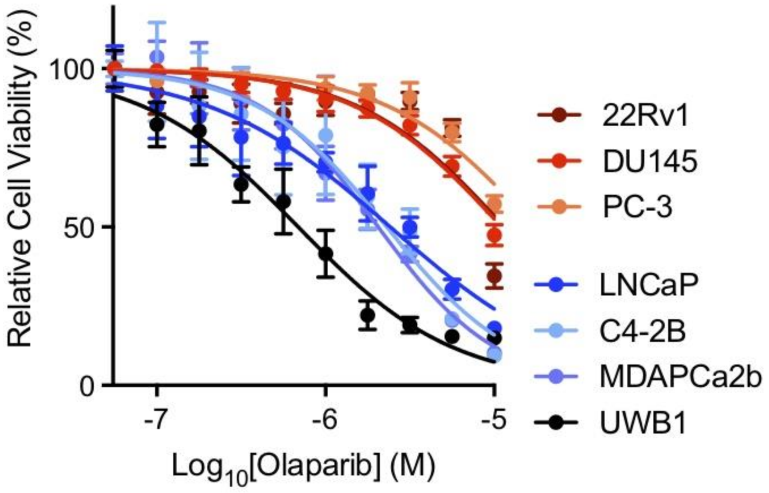


**Supplementary Fig. 1 The response of human cancer cell lines to PARP inhibitor olaparib.** Dose response curves after treatment with olaparib as indicated for 7 days in six human prostate cancer (PCa) cell lines (LNCaP, C4-2B, MDAPCa2b, 22Rv1 DU145 and PC-3) and one BRCA1-null human ovarian cancer cell line UWB1. Error bars represent SD, n = 6.

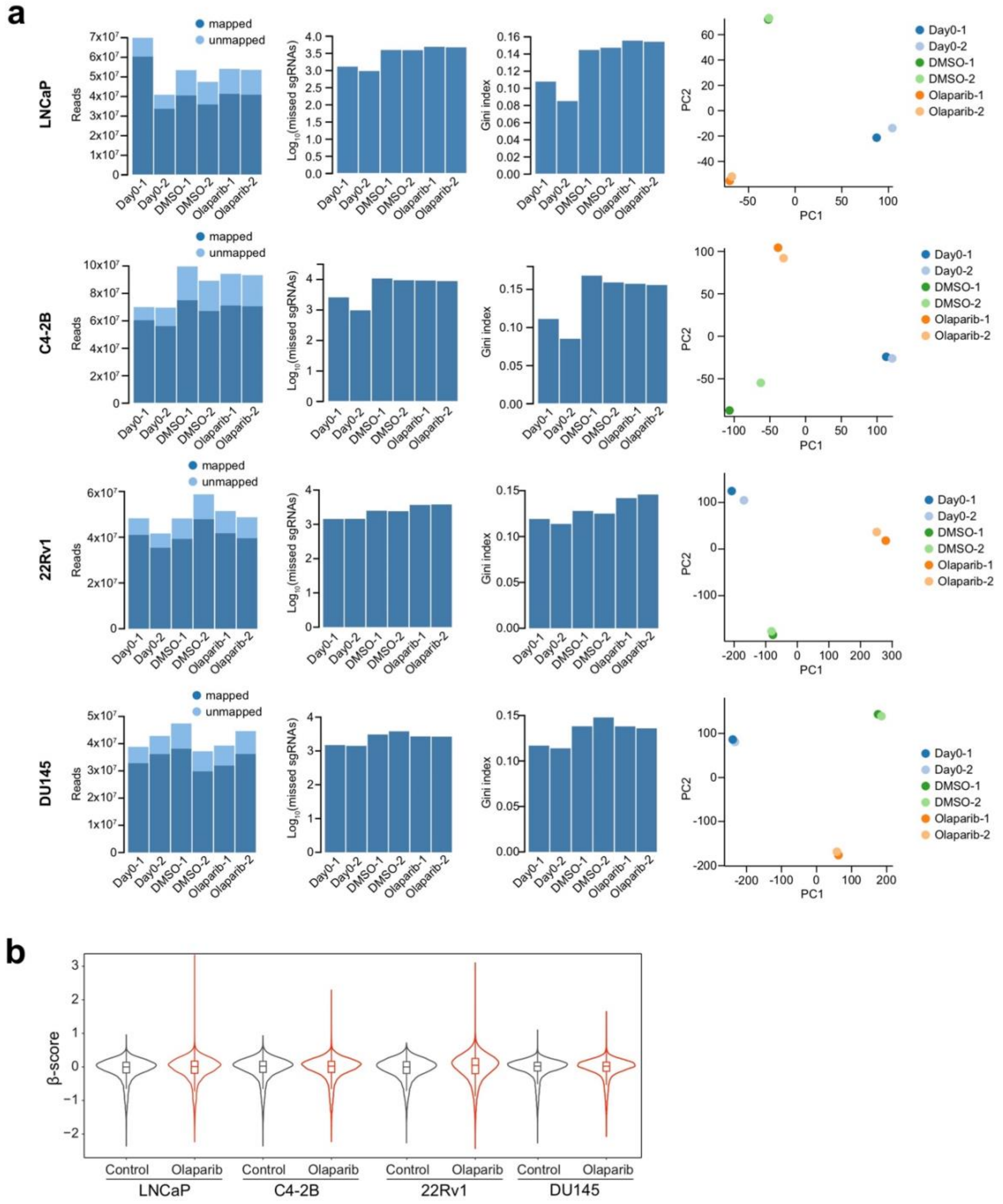


**Supplementary Fig. 2 The quality control measurements of CRISPR screens in four PCa cell lines. a,** Read counts of mapped and unmapped sgRNAs, number of missing sgRNAs, Gini index which is the measurement of read evenness within samples, and principal component analysis in each cell line are shown. **b,** Violin plots indicating the distribution of β-scores in each screen. Solid lines and boxes inside violin indicate median and interquartile range (IQR), respectively. Whiskers inside violin indicate the range of 1.5 x IQR.

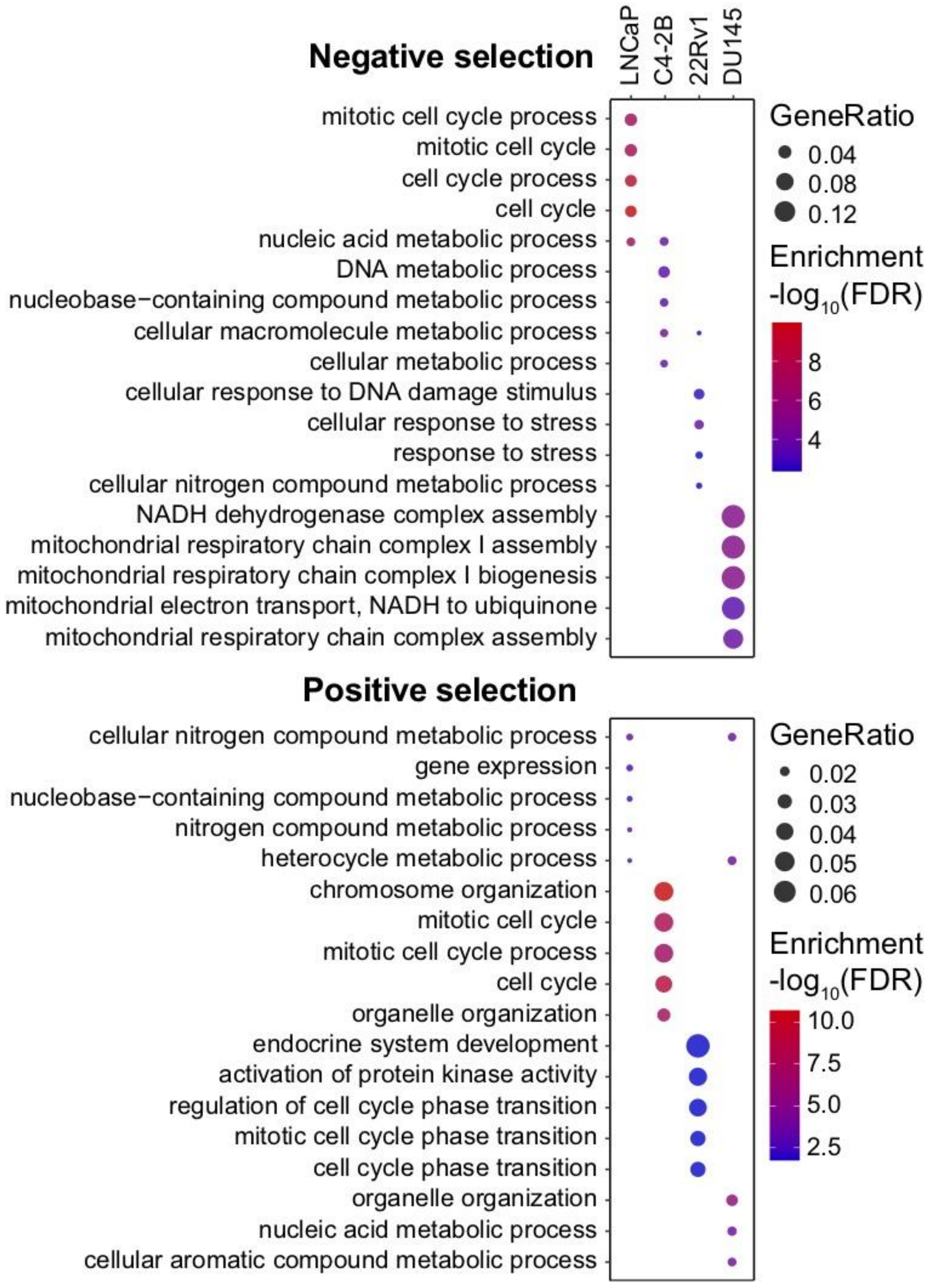


**Supplementary Fig. 3 Gene Ontology (GO) analysis of unique genes identified in each CRISPR screen after exclusion of common hits.** Top GO terms enriched in unique negatively selected genes in each cell line after exclusion of common hits (upper panel: LNCaP, n = 167; C4-2B, n = 195; 22Rv1, n = 129; DU145, n = 180). Top GO terms enriched in unique positively selected genes in each cell line after exclusion of common hits (lower panel: LNCaP, n = 211; C4-2B, n = 192; 22Rv1, n = 272; DU145, n = 236).

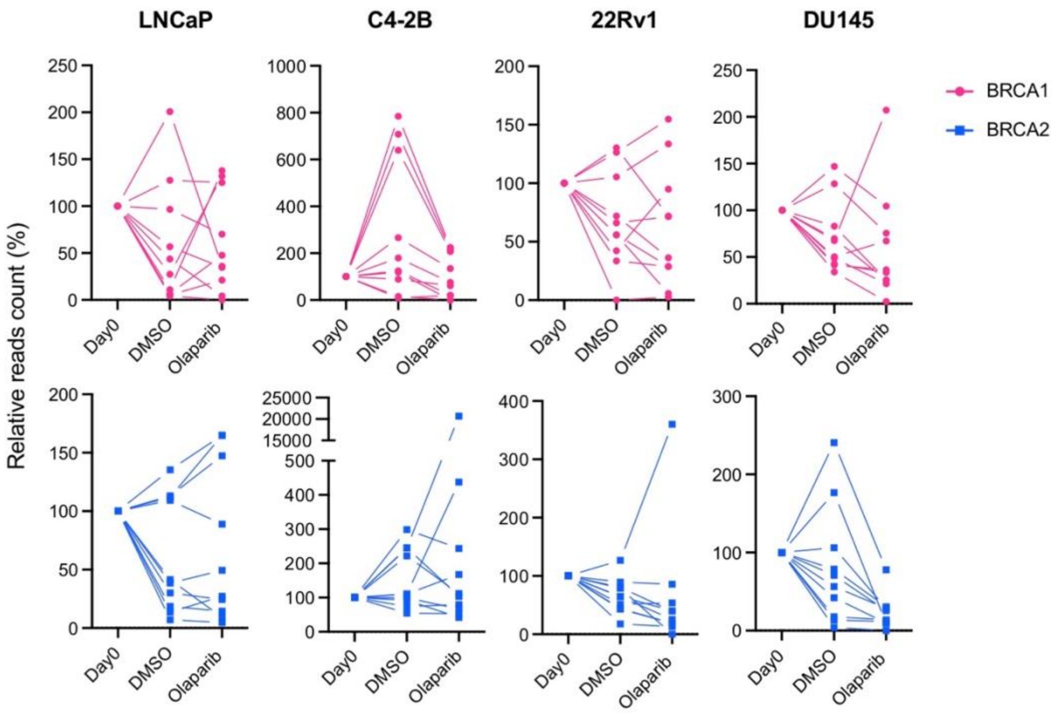


**Supplementary Fig. 4 BRCA1 and BRCA2 function as fitness genes.** Reads counts of BRCA1 and BRCA2 sgRNAs in CRISPR screen samples from day 0 and day 28 with DMSO or olaparib treatment. The read count at day 0 is defined as 100.

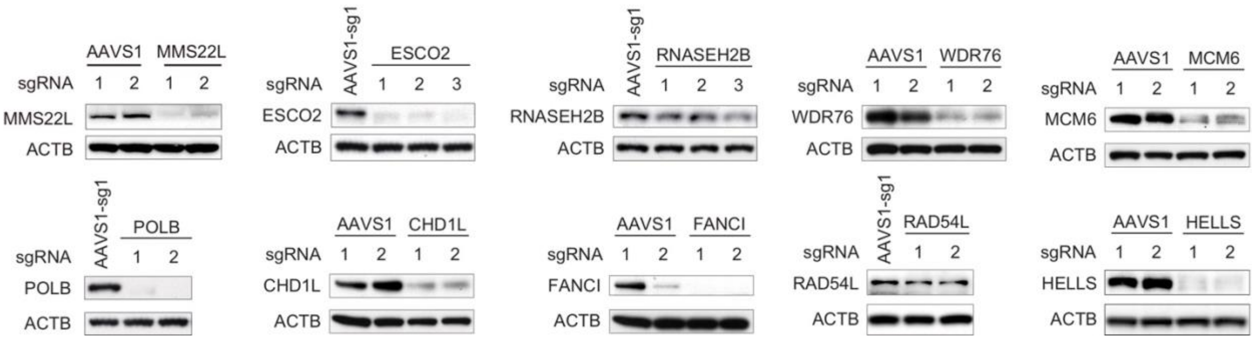


**Supplementary Fig. 5 Validation of gene knockout (KO).** Immunoblot analyses of the protein level of each gene as indicated after CRISPR/Cas9 KO in C4-2B cells. ACTB (β-actin) is a loading control.

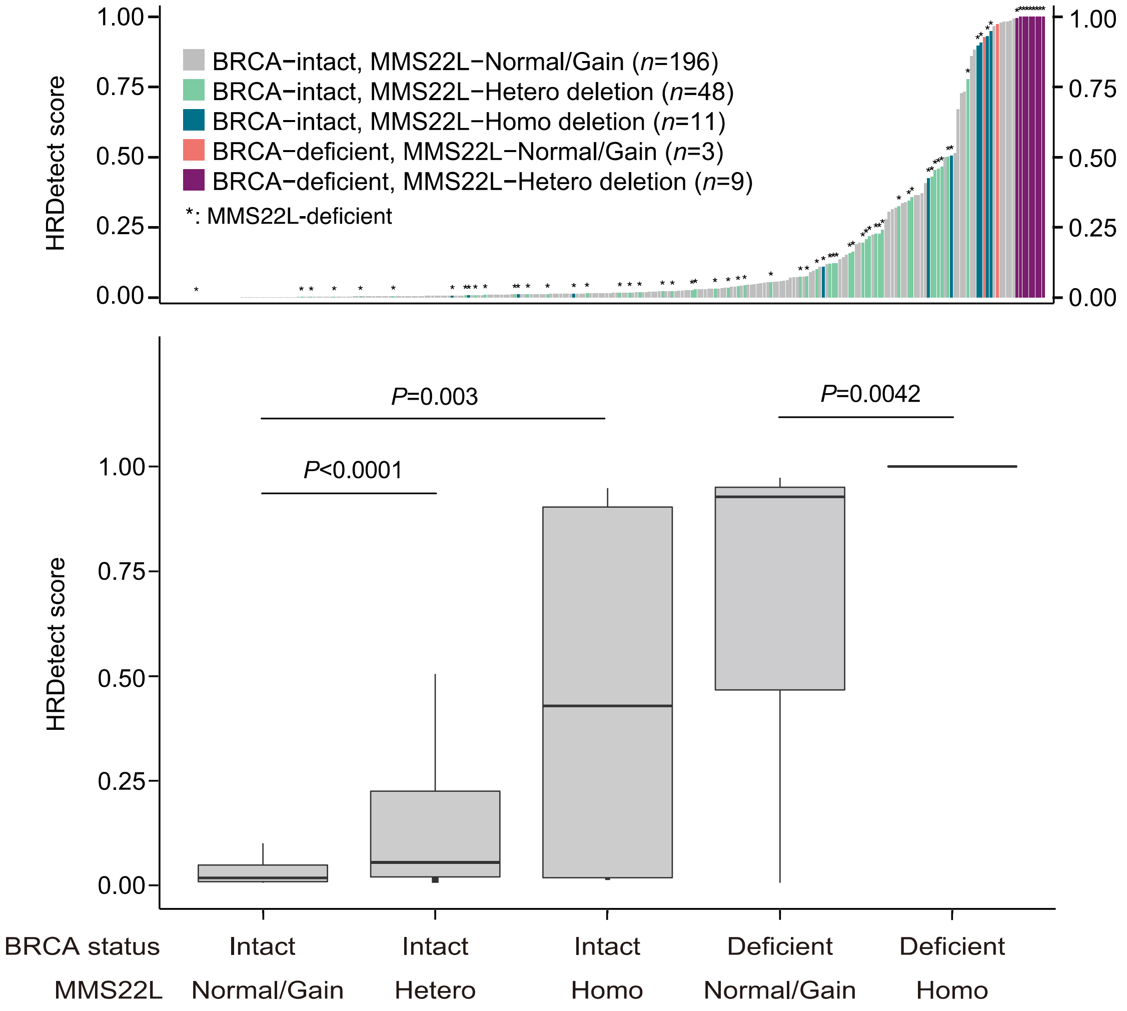


**Supplementary Fig. 6 HRDetect scores in prostate tumors with BRCA and/or MMS22L alterations.** Ranked HRDetect scores (upper panel) in prostate tumors with BRCA and/or MMS22L genomic alterations as indicated. Comparison of HRDetect scores (lower panel) between patients groups classified based BRCA and MMS22L status. HRDetect score is calculated using whole-genome sequencing data as previously described (see **Fig. 4e**). The *p*-values were determined using unpaired t-test.

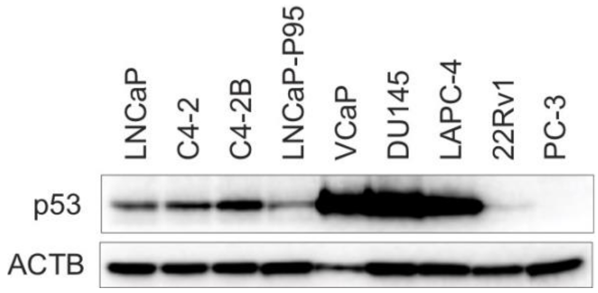


**Supplementary Fig. 7 p53 protein expression in PCa cell lines.** Immunoblot analysis of endogenous p53 protein expression in PCa cell lines as indicated. ACTB (β-actin) is a loading control.

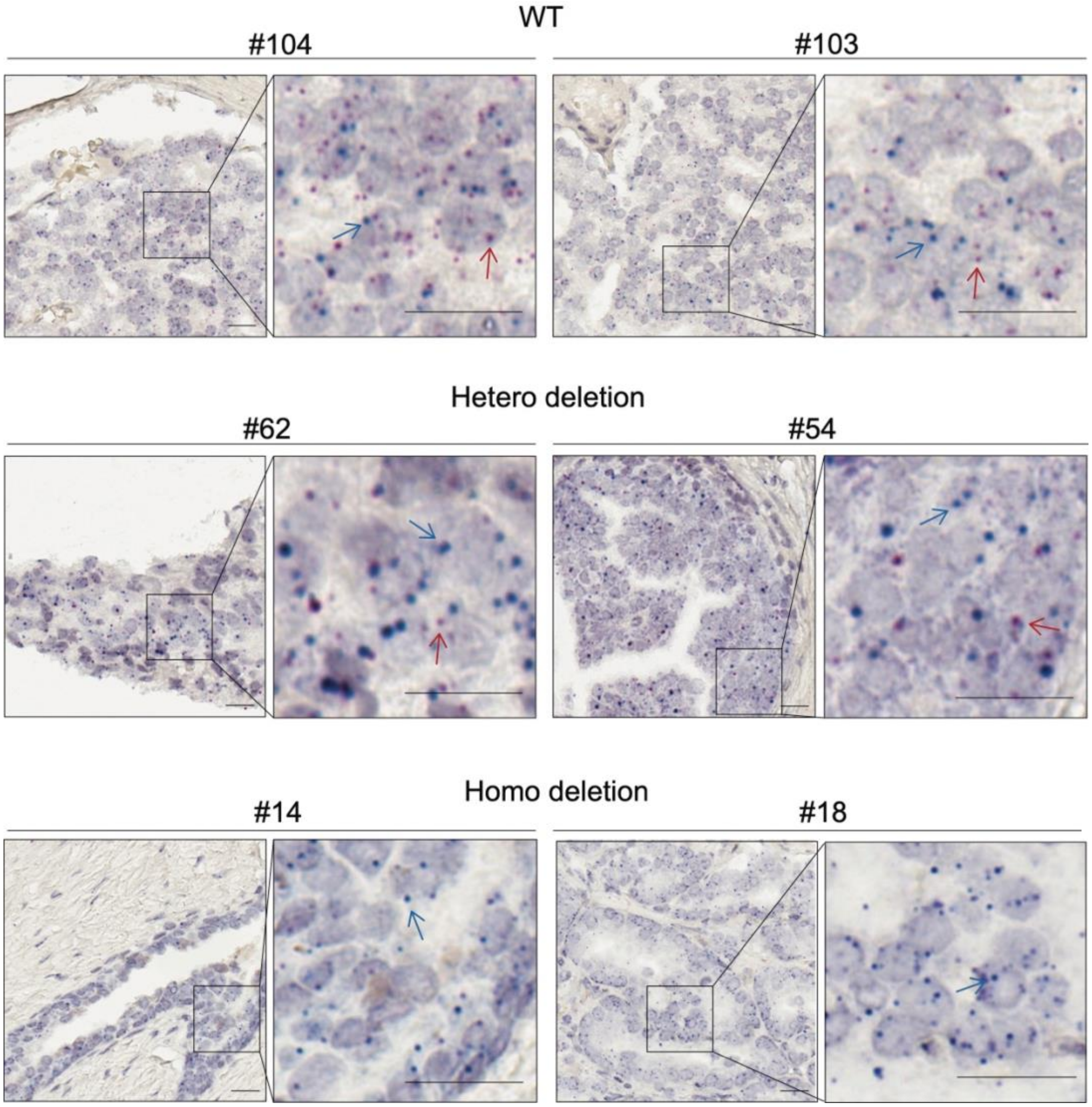


**Supplementary Fig. 8 DNAscope Assay on PCa tissue microarray (TMA).** Two representative MMS22L DNAscope images from MMS22L wild-type, heterozygous deletion, and homozygous deletion are presented, respectively. Red signals (red arrow) indicate a probe targeting the MMS22L gene on chromosome 6q. Blue signals (blue arrow) indicate a chromosome enumeration control probe targeting the centromeric region of chromosome 6p (CEP6p). The number of red and blue dots were counted and the ratio of red/blue was calculated. WT, red/blue > 0.5; Hetero deletion, red/blue between 0.1-0.5; Homo deletion, red/blue < 0.1. Scale bar = 20μm.

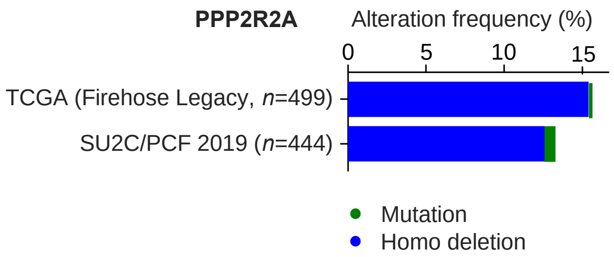


**Supplementary Fig. 9 PPP2R2A genomic alterations in PCa.** Frequency of PPP2R2A mutation and homozygous (Homo) deletion in the TCGA (Firehose Legacy, n = 499) and SU2C/PCF (n = 444) cohorts.

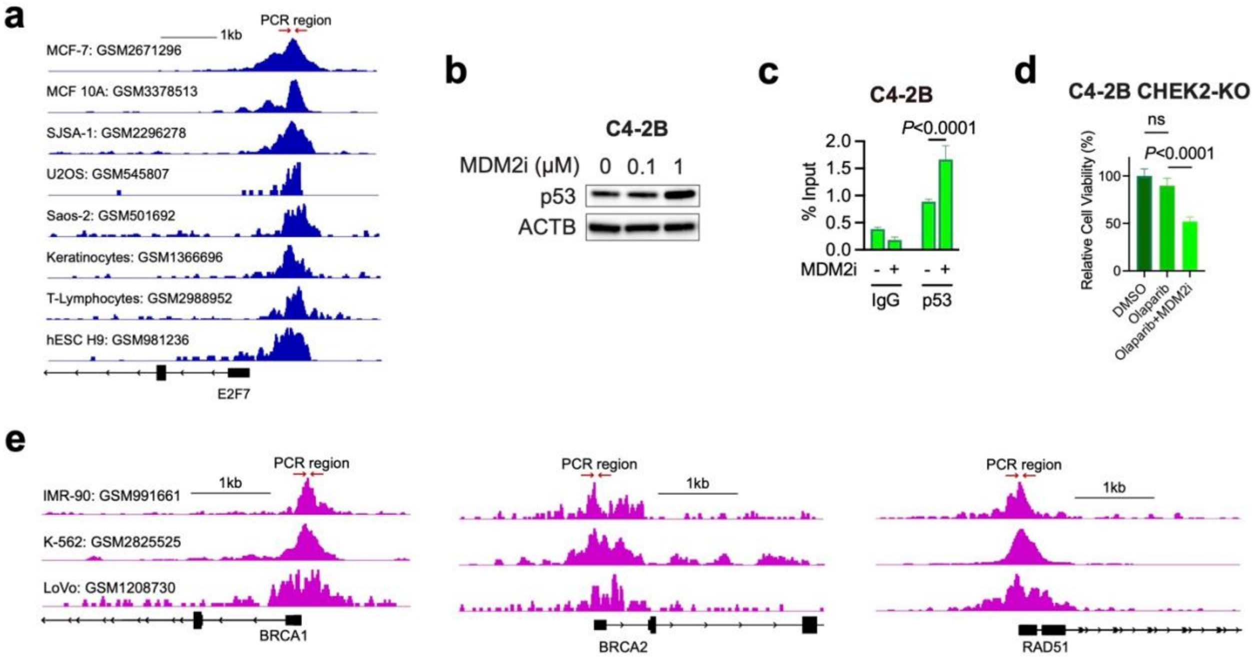


**Supplementary Fig. 10 p53- and E2F7-mediated gene regulation. a,** Genome browser view of p53 ChIP-seq signals at the E2F7 promoter in the indicated cell lines. The ChIP-seq data were obtained from publicly available datasets as indicated. Red arrows indicate the genomic region used for ChIP-qPCR validation in **Fig. 7d**. **b,** Immunoblot analysis of p53 in C4-2B cells after treatment with MDM2 inhibitor (MDM2i) nutlin as indicated. **c,** p53 ChIP-qPCR was performed at the E2F7 promoter region in C4-2B cells after treatment with nutlin (1 μM) or vehicle for 24 h. p53 occupancy was significantly increased at the E2F7 promoter after MDM2 inhibition. **d,** Cellular viability after treatment with olaparib (5 μM) in the presence or absence of MDM2i nutlin (1 μM) in CHEK2-KO C4-2B cells. Error bars represent SD, n = 6. The *p*-value was determined using unpaired *t*-test. **e,** Genome browser view of E2F7 ChIP-seq signals at the promote regions of BRCA1, BRCA2, and RAD51 genes in the indicated cell lines. The ChIP-seq data were obtained from publicly available datasets as indicated. Red arrows indicate the genomic region used for ChIP-qPCR validation in **Fig. 7f**.

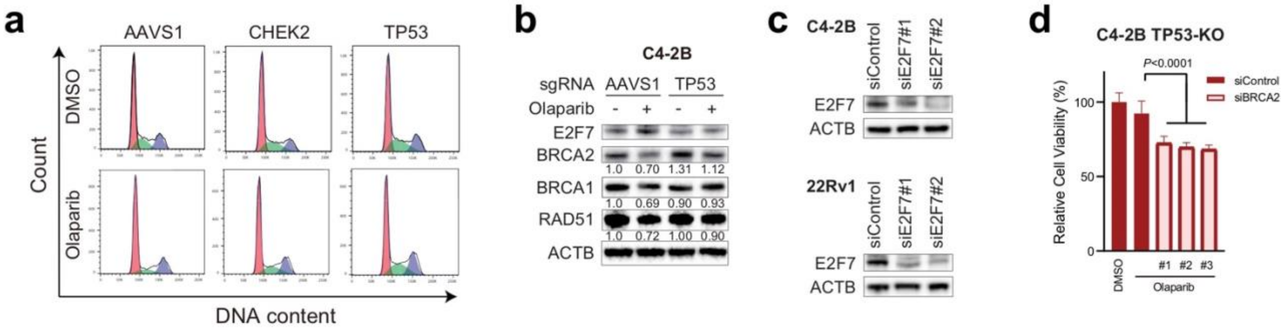


**Supplementary Fig. 11 Loss of TP53 increases HRR gene expression. a,** Cell cycle analysis in AAVS1 control, CHEK2-KO, and TP53-KO cells after treatment with DMSO or olaparib (5 μM) for 72 hours**. b,** Immunoblot analysis of the indicated proteins in AAVS1 control and TP53-KO C4-2B cells after treatment with olaparib (1 μM) or vehicle. The protein levels of BRCA1, BRCA2, and RAD51 genes decreased after olaparib treatment in TP53-intact AAVS1 control C4-2B cells but remained at a high level in TP53-KO C4-2B cells. The integrated optical density (IOD) values of the indicated proteins normalized by ACTB IOD are shown. **c,** Immunoblot analyses of E2F7 protein levels in C4-2B and 22Rv1 cells transfected with siRNAs against E2F7 or negative control. **d,** Cellular viability after olaparib (1 μM) treatment for 7 days in TP53-KO C4-2B cells transfected with siRNAs against BRCA2 or negative control. The *p*-value was determined by unpaired t-test. Error bars represent SD, n = 6.

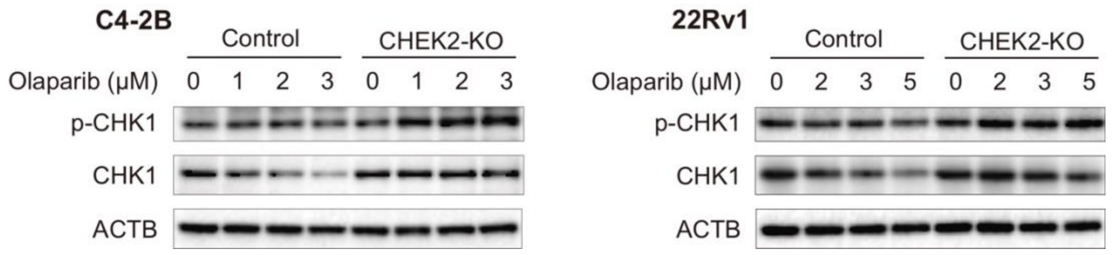


**Supplementary Fig. 12 Olaparib treatment induces ATR activity.** Immunoblot analyses of p-CHK1 and CHK1 in AAVS1 control and CHEK2-KO C4-2B and 22Rv1 cells after olaparib treatment as indicated.

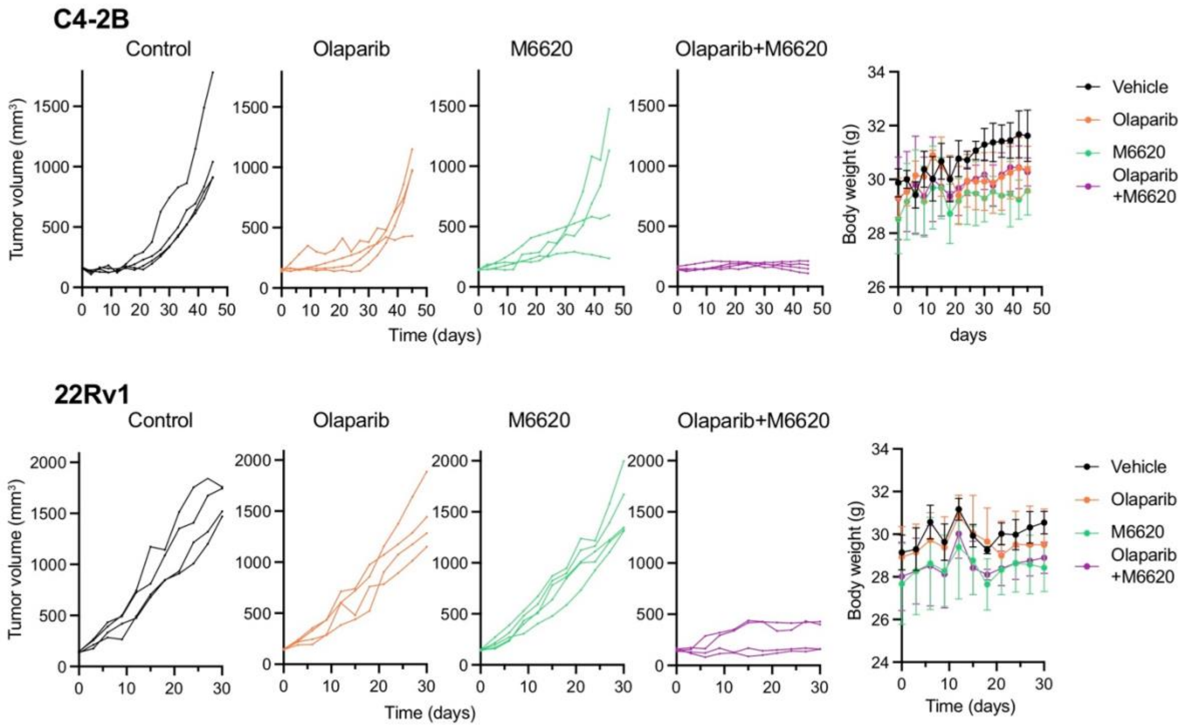


**Supplementary Fig. 13 Combination treatment with olaparib and M6620 in CHEK2-KO xenograft models.** Tumor growth of CHEK2-KO C4-2B and 22Rv1 cells in each mouse treated with the indicated agents and mean body weight of the mice in each group are shown.
